# Understanding transformation tolerant visual object representations in the human brain and convolutional neural networks

**DOI:** 10.1101/2020.08.11.246934

**Authors:** Yaoda Xu, Maryam Vaziri-Pashkam

## Abstract

Forming transformation-tolerant object representations is critical to high-level primate vision. Despite its significance, many details of tolerance in the human brain remain unknown. Likewise, despite the ability of convolutional neural networks (CNNs) to exhibit human-like object categorization performance, whether CNNs form tolerance similar to that of the human brain is unknown. Here we provide the first comprehensive documentation and comparison of three tolerance measures in the human brain and CNNs. We measured fMRI responses from human ventral visual areas to real-world objects across both Euclidean and non-Euclidean feature changes. In single fMRI voxels in higher visual areas, we observed robust object response rank-order preservation across feature changes. This is indicative of functional smoothness in tolerance at the fMRI meso-scale level that has never been reported before. At the voxel population level, we found highly consistent object representational structure across feature changes towards the end of ventral processing. Rank-order preservation, consistency, and a third tolerance measure, cross-decoding success (i.e., a linear classifier’s ability to generalize performance across feature changes) showed an overall tight coupling. These tolerance measures were lower for Euclidean than non-Euclidean feature changes in lower visual areas, but increased over the course of ventral processing in most cases. These characteristics of tolerance, however, were absent in eight CNNs pretrained with ImageNet images with varying network architecture, depth, the presence/absence of recurrent processing, or whether a network was pretrained with the original or stylized ImageNet images that encouraged shape processing. Most notably, CNNs do not show increased representational consistency across feature changes at the higher layers. CNNs thus do not appear to develop the same kind of tolerance as the human brain over the course of visual processing.

**Significant Statement:** Perceiving object identity among changes in non-identity features and forming transformation-tolerant object representations is essential to high-level primate vision. Here we provide a comprehensive documentation and comparison of three tolerance measures between the human brain and CNNs pretrained for object classification. While all three measures show increased tolerance in the human brain across four types of feature changes towards the end of ventral visual processing, CNNs fail to develop the same kind of tolerance with visual processing.

## Introduction

One of the hallmarks of primate high-level vision is its ability to extract object identity features among changes in non-identity features and form transformation-tolerant object representations (DiCarlo & Cox, 2007; DiCarlo et al., 2012; Tacchetti et al., 2018). This allows us to rapidly recognize an object under different viewing conditions. Computationally, achieving tolerance reduces the complexity of learning by requiring fewer training examples and improves generalization to objects and categories not included in training (Tacchetti et al., 2018). Despite its significance, many details of tolerance in the human brain remain unknown, such as its functional smoothness, representational consistency across a feature change, and the relationship among the different tolerance measures. Recent developments in hierarchical convolutional neural networks (CNNs) have generate the excitement that perhaps the algorithms essential to high-level primate vision would automatically emerge in CNNs to provide us with a shortcut to understand and model high-level vision. Whether CNNs form tolerance similar to that of the primate brain, however, has not been thoroughly investigated. Here we provide the first comprehensive documentation of tolerance in the human brain and CNNs, and a comparison between the two. We measured fMRI responses from the human ventral visual areas to real-world objects across four types of nonidentity feature changes including both Euclidean and non-Euclidean features. We examined responses both at the single voxel level and the voxel population level and documented as well as compared three tolerance measures, including object response rank-order preservation, cross-decoding success and representational consistency. We also documented the same tolerance measures in 8 CNNs pretrained with ImageNet images (Deng et al., 2009) with varying network architecture, depth, and the presence/absence of recurrent processing, as well as a network pretrained with either the original or stylized ImageNet images that encouraged shape processing (Geirhos et al., 2019).

At the single neuron level, tolerance is reflected in macaque IT neurons’ ability to preserve the rank order of the neuronal firing rate to different objects across a nonidentity feature change even when the absolute responses rescale with the change (Schwartz et al., 1983; Tovee et al., 1994; Ito et al., 1995; DiCarlo & Manusell, 2003; Brincat & Connor, 2004; DiCarlo and Cox, 2007; Li et al., 2009; Murty & Arun, 2017; see Figure 1a and 1b for a schematic illustration). At the population level, tolerance is seen as the ability of a group of neurons in macaque IT or fMRI voxels in human occipito-temporal cortex (OTC) to decode object identity across nonidentity feature changes (e.g., testing if a linear classifier trained to differentiate objects at one position is successful at doing so at an untrained position; Hung et al., 2005; Rust & DiCarlo, 2010; Cichy et al., 2011; Vaziri-Pashkam & Xu, 2019; Vaziri-Pashkam et al., 2019; Taylor & Xu, 2022; see also Ward et al., 2018; Mocz et al., 2021). Neuronal recording and simulation further show that rank-order preservation at the single neuron level and cross-decoding success at the population level are closely correlated (Li et al., 2009). Such a correlation predicts that rank-order is likely preserved in individual fMRI voxels to support cross-decoding at the voxel population level.

**Figure 1.**
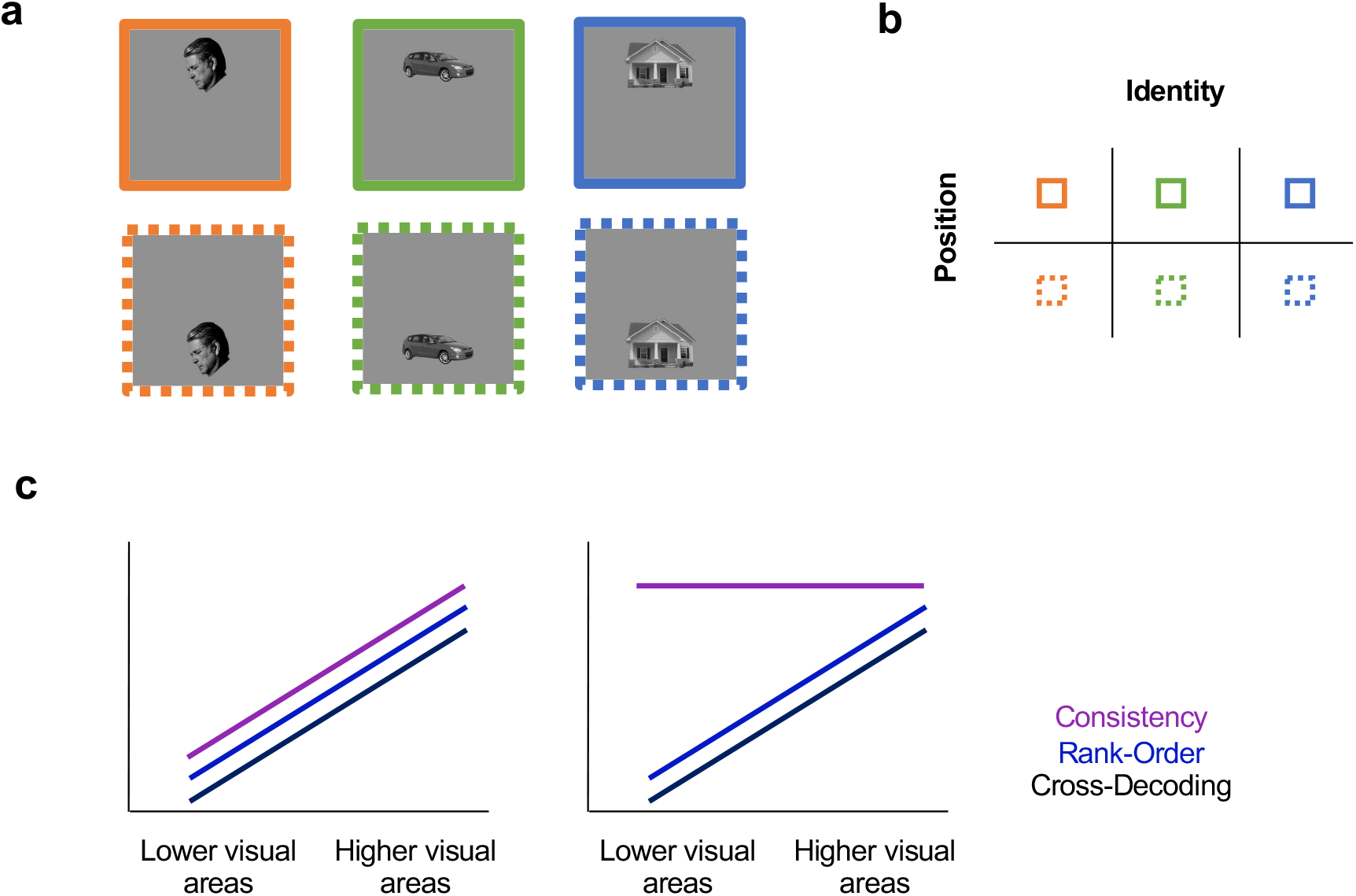
Illustrations of the object representational space and possible tolerance measures to three objects appearing at two positions during visual processing. **a.** Three example objects appearing at two different spatial positions. **b.** Orthogonal object identity and position representations at higher visual areas to allow independent readout of these two types of information. In such a representational space, the similarity among a set of objects is well preserved across the position change. **c.** The coupling of representational consistency with response rank-order preservation and cross-decoding success. Left, a tight coupling among the three tolerance measures, with values going up from lower to higher visual areas; right, a decoupling of consistency from the two other measures, with consistency being relatively constant over the course of visual processing if there is uniformity in visual representation in visual areas.

Given that each fMRI voxel contains around a million neurons (Huettel et al., 2009), rank-order preservation at the fMRI voxel meso-scale level is not obvious and has never been examined before. Past neurophysiological studies have only reported rank-order preservation in isolated IT neurons. Whether there is functional smoothness in high-level vision to enable rank-order preservation at the single fMRI voxel level, however, is unknown. Can we see tolerance in individual fMRI voxels? If so, how does it emerge over the course of ventral visual processing? Is there a close correspondence between rank-order preservation and cross-decoding success in fMRI measures?

Our connection with the world involves not only identifying objects but also interacting with them. To do so, nonidentity information about the objects, such as position and size, also needs to be represented during visual processing. Both object identity and nonidentity features can be independently read out across changes in the other feature in primate higher visual areas, with these two types of features represented in a largely distributed and overlapping manner in single neurons (e.g., Hung et al., 2005; Schwarzlose et al., 2008; Sayres & Grill-Spector, 2008; Carlson et al., 2011; Cichy et al., 2011; Hong et al., 2016; Vaziri-Pashkam et al., 2019; Mocz et al., 2021). The independent readout of identity and nonidentity features critically requires orthogonal feature representations and representational consistency in the feature space, such that the similarity among a set of objects is well preserved across a non-identity feature change (see Figures 1a and 1b for a schematic illustration). Representational consistency at the population level, despite it being necessary to achieve tolerance, has never been directly documented. While cross-decoding success has been used to evaluate tolerance, it emphasizes the success of representation generalization across feature changes. It is not sensitive to changes in the representational space, so long as objects are still on the correct side of the decoding decision boundary. Consequently, cross-decoding does not accurately depict changes in the representational space after a nonidentity feature change.

Consistency may be closely associated with rank-order preservation at the single neuron level. If so, given the increase of the latter with visual processing, consistency is expected to increase as well from lower to higher ventral visual areas (Figure 1c left). Consistency, however, may be dissociable from rank-order preservation. For example, due to the small neuronal receptive field size of early visual areas and the presence of retinotopy, largely nonoverlapping neurons would represent an object at different positions and sizes, leading to a lack of rank-order preservation and a failure of cross-decoding across these feature changes. However, as long as these different neuronal populations represent objects in a similar way, consistency can still be maintained. Consistency may thus be decoupled from rank-order preservation and cross-decoding success if there is uniformity in visual representation in visual areas (Figure 1c right).

Here we examined these three tolerance measures (i.e., rank-order preservation, cross-decoding success and representational consistency) across changes in two commonly studied Euclidean features (i.e., position and size) and two rarely studied non-Euclidean features (i.e., image stats and SF). We manipulated the image stats and, from the original real-world images, generated controlled images in an effort to normalize and equalize the spectrum, histogram, and intensity of the images across the different categories (Willenbockel et al., 2010). This ensured that object identity representation in lower visual areas would reflect the representation of identity and not low-level image differences among the objects from different categories. As observers had no problem recognizing objects undergone such a change, this provided us with a unique opportunity to examine how tolerance to such image stats change would develop over the course of ventral visual processing. Besides changes to image stats, we can easily recognize objects both near and far despite large changes in the spatial frequency (SF) content of an image. We also have the remarkable ability of recognizing line-drawing objects although they do not exist in natural vision. Tolerance to SF change, to our knowledge, has never been studied in detail. By studying tolerance to both Euclidean and non-Euclidean feature changes, we can develop a better understanding of whether the development of tolerance to these features follow a similar or different profile from lower to higher human ventral visual areas.

Recent hierarchical convolutional neural networks (CNNs) have achieved human-like object categorization performance and are able to identify objects across a variety of identity preserving image transformations (Yamins & Dicarlo, 2016; Kheradpisheh et al., 2016; Kriegeskorte, 2015; Serre, 2019; Han et al., 2020; Blything et al., 2021). Moreover, both shallow and deep CNNs can successfully decode objects across changes such as position, size, and viewpoint/rotation (Kheradpisheh et al., 2016). This has led to the thinking that CNNs form transformation-tolerant object representations in their final stages of visual processing similar to those seen in the primate brain (Hong et al., 2016; Yamins & Dicarlo, 2016; Tacchetti et al., 2018). Although CNNs are fully image computable and accessible, they are at the same time extremely complex models with millions or even hundreds of millions of free parameters. The general operating principles at the algorithmic level (Marr, 1982) that enable CNNs’ success in object categorization, however, remain poorly understood (Kay, 2018). Can we see rank-order preservation in individual CNN units? What about representational consistency? Can we find a brain-like relationship among rank-order preservation, cross-decoding success and consistency in CNNs over the course of visual processing?

To anticipate, in human higher visual areas, we found robust rank-order preservation in individual fMRI voxels, indicative of functional smoothness of tolerance, and highly consistent object representational structure across nonidentity feature changes. Moreover, rank-order preservation, cross-decoding success and consistency were tightly correlated. These tolerance measures were lower for Euclidean than non-Euclidean feature changes in lower visual areas, but all increased over the course of ventral processing in most cases. Such a relationship, however, was absent in the 8 CNNs tested regardless of differences in architecture, depth, the presence/absence of recurrent processing, or training. CNNs thus do not appear to develop the same kind of tolerance as the human brain over the course of visual processing.

## Results

In four fMRI experiments, human participants viewed blocks of sequentially presented object images. Each image block contained different exemplars from the same object category. A total of eight real-world object categories were used, and they were bodies, cars, cats, chairs, elephants, faces, houses, and scissors (Vaziri-Pashkam & Xu, 2019; see Figure 2a). These object images were shown in two types of Euclidean feature changes (Figure 2b): position (top vs bottom) and size (small vs large), and two types of non-Euclidean feature changes (Figure 2b): image stats (original vs controlled) and SF (high vs low). To ensure that object identity representation in lower brain regions would reflect the representation of identity and not low-level image differences among the different categories, controlled images were also used for position and size changes. To increase signal to noise ratio (SNR), we examine the averaged response from multiple images from the same object category rather than the response of a single image. Previous research has shown similar category and exemplar response profiles in macaque IT and human lateral occipital cortex with more robust responses obtained for categories than individual exemplars due to an increase in SNR (Hung et al., 2005; Cichy et al., 2011). Rajalingham, et al. (2018) further reported better behavior-CNN correspondence at the category but not at the individual exemplar level. Thus, obtaining fMRI responses at the category level, rather than at the exemplar level, could only increase our chance of finding a close brain-CNN correspondence if it exists.

**Figure 2.**
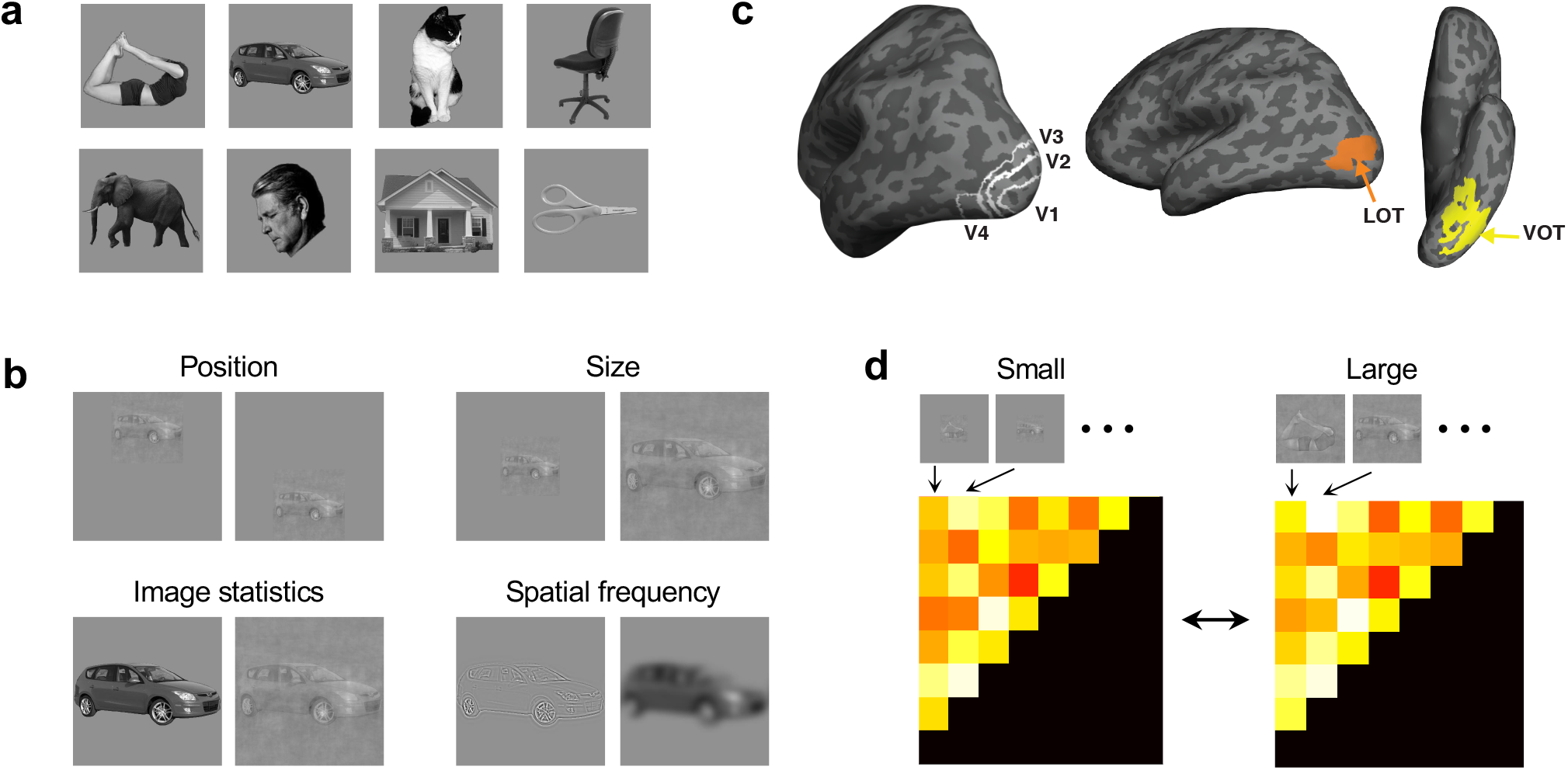
Stimuli used, brain regions examined, and the consistency analysis applied in the study. **a.** The eight real-world object categories used with an example image shown for each category. Ten different exemplars varying in identity, viewpoint/orientation and pose/expression (for body, cat, elephant, face and scissor only) were used for each category. **b.** Example images of the two Euclidean and two non-Euclidean feature changes examined. The Euclidean feature changes included changes in position (top vs bottom) and size (small vs large). The non-Euclidean feature changes included changes in image stats (original vs controlled) and the SF content of an image (high vs low). Controlled images were created by normalizing and equalizing the spectrum, histogram, and intensity of the images across the different categories. To ensure that object identity representation in lower visual areas reflected the representation of identity and not low-level differences among the images of the different categories, controlled images were used for both the position and size changes.**c.** The brain regions examined. They included topographically defined early visual areas V1 to V4 and functionally defined higher object processing regions LOT and VOT. **d.** Measuring object representational space consistency across a nonidentity feature change (using size change as an example). For a given brain region or a sampled CNN layer, a representational dissimilarity matrix was first constructed by computing all pairwise Euclidean distances of fMRI response patterns or CNN output for all the object categories at one value of a nonidentity feature. The off-diagonal elements of this matrix were then used to construct a representational dissimilarity vector. The dissimilarity vectors from two values of a nonidentity feature were correlated to evaluate representational space consistency across the feature change, with a high correlation indicating a high consistency.

We examined fMRI responses from independently defined human early visual areas V1 to V4 and higher visual object processing regions LOT and VOT (Figure 2c). Reponses in LOT and VOT have been shown to correlate with successful visual object detection and identification (Grill-Spector et al. 2000; Williams et al., 2007) and their lesions have been linked to visual object agnosia (Goodale et al.,1991; Farah, 2004). These two regions have been argued to be the homologue of the macaque IT (Orban et al., 2004). For a given brain region, fMRI response patterns were extracted for each category for each type of nonidentity feature changes.

### Single fMRI voxel response rank order preservation for objects across nonidentity feature changes

To evaluate rank-order preservation across a nonidentity feature change, using Spearman rank correlation, we correlated a voxel’s responses to all the object categories shown at one value of the nonidentity feature to those at the other value. To account for region-specific noise and ensure valid comparisons across brain regions, these correlations were corrected by the reliability of each brain region using a split-half measure (see Methods; Figure 3b). In lower visual areas, while the corrected rank-order correlations were below .37 for the two Euclidean feature changes, they were above .53 for the two non-Euclidean feature changes. In fact, the corrected rank-order correlations were no greater than 0 in areas V1 and V2 for position change *(ts* < 1.50, *ps* > .11; all one tailed and corrected for multiple comparisons for the six brain regions included for each nonidentity feature change; one-tailed t tests were used here to test for the difference in a specific direction; this applies to all subsequent t tests of the same type); for all others, they were significantly above 0 (*ts* > 2.11, *ps* < .040). At higher visual areas, rank-order was fairly well preserved, such that, for all four feature changes, at least one higher visual area showed a corrected rank-order correlation either not significantly lower than 1 or only marginally lower than 1 (for position, LOT, *t(6)* = 1.69, *p* = .071; for size, V4, *t(6)* = 1.92, *p* = .078, LOT, *t(6)* = 1.71, *p* = .083, and VOT*, t(6)* = 1.57, *p* = .084; for image stats, V2, *t(5)* = 2.71, *p* = .064, V3, *t(5)* = 2.35, *p* = .066, V4, LOT and VOT, *t* < 1.75, *ps* > .11; for SF, LOT, *t(10)* = 1.48, *p* = .086; and for all others, *ts* > 1.99, *ps* < .047). Further analyses revealed a positive relationship between these rank-order correlation coefficients and the rank order of the visual areas for all feature changes (*rs* > .45, *ts* > 3.19, *ps* < .0079; one-tailed and corrected for the four nonidentity features included, one-tailed t tests were used here to assess the correlation in a specific direction; this applies to all subsequent t tests of the same type). Direct comparisons further confirmed that, for all four feature changes, the average rank-order correlation coefficients of V1 and V2 were lower than those of LOT and VOT (*ts* > 3.29, *ps* < .0055).

**Figure 3.**
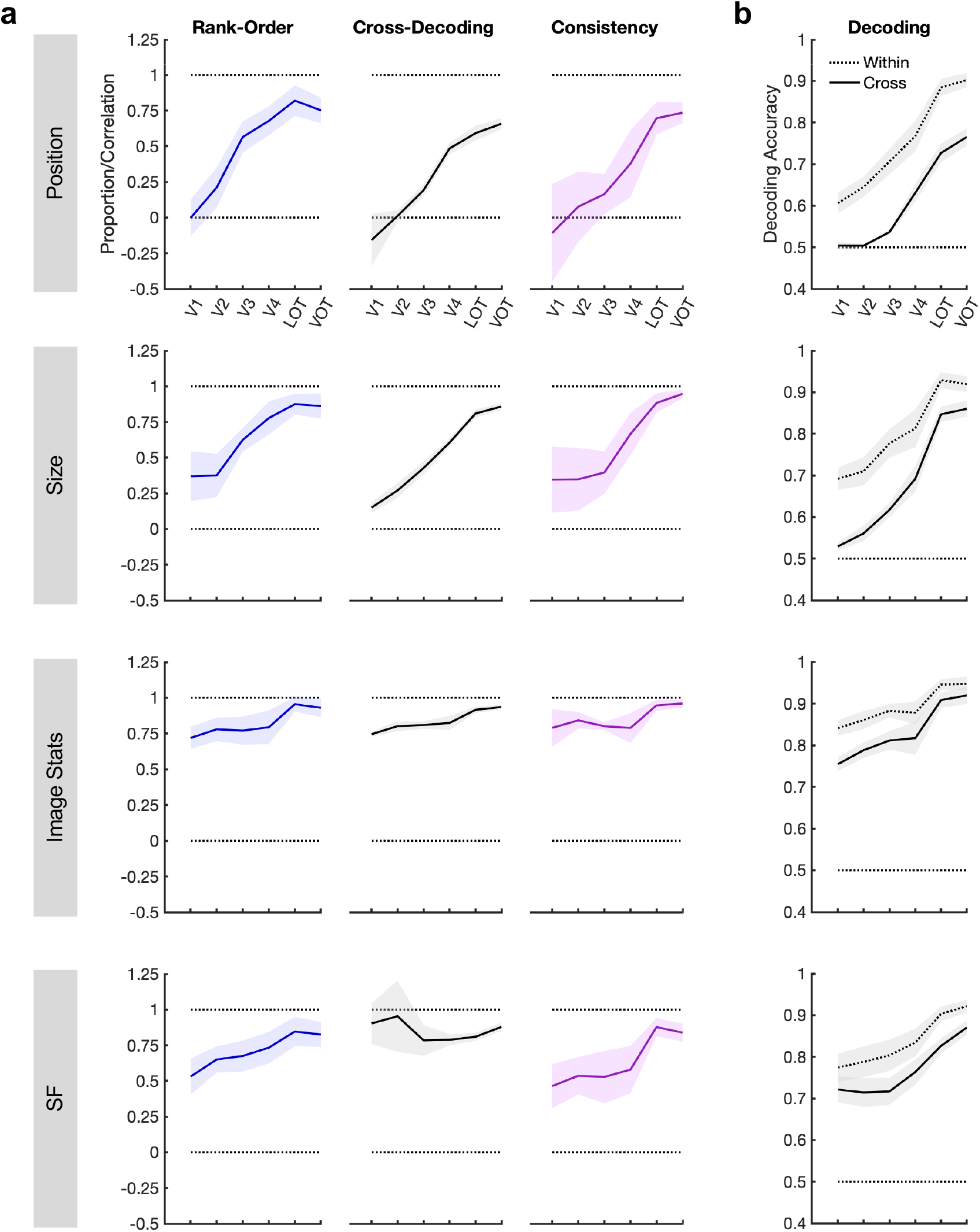
Tolerance measures from human ventral visual areas V1 to V4, LOT and VOT across the two Euclidean (position and size) and the two non-Euclidean (image stats and SF) feature changes. **a.** Rank-order preservation, cross-decoding success, and consistency across the brain areas. Rank-order preservation was measured as the Spearman correlation of a voxel’s responses to all the object categories shown at one value of a nonidentity feature to those at the other value, corrected by the reliability of each brain region (see Methods). Cross-decoding success was measured as a ratio and obtained by first subtracting 0.5 from the raw within- and cross-decoding accuracies (see b) and then taking the ratio of the two resulting values. Consistency was calculated as the representational structure correlation across a nonidentity feature change (see Figure 2d), corrected by the reliability of each brain region (see Methods). **b.** Object category decoding accuracy with no feature change (within-decoding) and across a nonidentity feature change (cross-decoding). The width of the ribbon represents the standard errors of the means.

At the single fMRI voxel level in human higher visual areas, we thus observed, for the first time, high rank-order preservation across nonidentity feature changes previously only reported for single neurons in macaque higher visual areas. Our examination of the entire ventral visual areas further revealed that, at lower visual areas, while rank-order preservation was low for the two Euclidean feature changes, it was already fairly high for the two non-Euclidean feature changes. That being said, rank-order preservation increased from lower to higher visual areas for both types of feature changes, showing that over the course of ventral processing, object response rank order become more aligned across a nonidentity feature change.

### Decoding objects within and across nonidentity feature changes in human ventral visual areas

What is the relationship between rank-order preservation at the single fMRI voxel level and cross-decoding success at the voxel population level? To answer this, we measured category decoding accuracy with no feature change (within-decoding; e.g., both training and testing were done for objects shown at the same position) and across a feature change (cross-decoding; e.g., training was done for objects at one position and testing was done for objects at another position) (Figure 3a). For all four types of feature changes, significant above chance within-decoding was found for all six brain regions examined (*ts* > 4.19, *ps* < .0029). Cross-decoding was at chance in areas V1 and V2 for position change (*ts* < .92, *ps* >.19), but was above chance for all other areas and feature changes (*ts* > 2.82, *ps* < .015). For all brain areas and changes examined, cross-decoding accuracy was lower than that of within-decoding (*ts* > 3.50, *ps* < .0076).

To combine the two decoding measures and to account for accuracy differences in within-decoding across the different brain areas and changes, we computed a decoding proportion score by first subtracting 0.5 from the within- and cross-decoding accuracies and then taking the ratio of the two resulting values. A proportion score of 1 would indicate equally good decoding performance within and across a feature change and a complete generalization of category representation across the nonidentity feature change, whereas a proportion score of 0 indicates a complete failure of such generalization (Figure 3b). For the two Euclidean features (i.e., position and size), the decoding proportion score started from below .16 in lower visual areas to rise above .65 in higher visual areas, whereas for the two non-Euclidean features (i.e., image stats and SF), the decoding proportion score was overall very high, ranging from .74 to .95. Detailed analyses showed that the decoding proportion scores were no greater from 0 in areas V1 and V2 for position change *(ts* < .28, *ps* > .47), but were above 0 for all other brain areas and feature changes (*ts* > 3.81, *ps* < .0037). Further analyses revealed a positive correlation between decoding proportion scores and the rank order of the visual areas for position, size and image stats changes (*rs* > .71, *ts* > 4.37, *ps* < .0036), but not for an SF change (*r* = .25*, t(10)* = 1.30, *p* = .11). Direct comparisons confirmed that, for position, size and image stats changes, the average proportion scores of V1 and V2 were lower than those of LOT and VOT (*ts* > 4.70, *ps* < .0035). This was not the case for the SF change (*t(10)* = .45, *p* = .67).

These results showed that category representation was robust for each value of the four nonidentity features in all six brain regions examined. With the exception of position change in V1 and V2, there were also significant, although incomplete, category representation generalization across changes. Such generalization appeared to be in general higher for changes involving non-Euclidean than Euclidean features. These generalizations increased from lower to higher ventral regions for changes involving position, size and image stats, but not SF. Overall, except for changes involving SF, cross-decoding success exhibited a similar response profile across visual areas as that of voxel rank-order preservation. The discrepancy seen with SF is likely due to decoding being a less sensitive measure of the representational space as discussed earlier. A quantitative comparison of these two measures is presented in a later analysis.

### Representational consistency of objects across nonidentity feature changes in human ventral visual areas

Is there representational consistency across nonidentity feature changes in higher visual areas? How does consistency evolve over the course of visual processing? To answer these questions, within each value of a given change, we first calculated pairwise Euclidean distances of fMRI response patterns for all the object categories to construct a category representational dissimilarity matrix (RDM, Kriegeskorte & Kievit, 2013, see Figure 2d and Methods). We then correlated these RDMs between the two values of each change using Spearman rank correlation. As before, to account for region-specific noise and ensure valid comparisons across brain regions, these correlations were corrected by the reliability of each brain region using a split-half measure before the results were compared across brain regions (see Methods). We used these corrected RDM correlations as our consistency measures (Figure 3b).

In lower visual areas, while consistency was below .35 for the two Euclidean feature changes, they were above .46 for the two non-Euclidean feature changes. In fact, consistency was no greater than 0 in areas V1 to V4 for position and in areas V1 and V2 for size (*ts* < 1.50*, ps* > .092); for all others, consistency was significantly above 0 (*ts* > 2.65, *ps* < .028). In higher visual areas, consistency was fairly high for both the Euclidean and non-Euclidean feature changes (all greater than .73), such that for size and for image stats in multiple visual areas it was either no lower or marginally lower than 1 (for size, LOT, *t(6)* = 1.70, *p* = .070, VOT, *t(6)* = 1.75, *p* = .079; for image stats, V2, *t(5)* = 2.74, *p* = .062, V4, *t(5)* = 2.02, *p* = .099, V1, LOT and VOT, *ts* <1.58, *ps* > .12; for all others, *ts* > 1.92, *ps* < .044). Further analyses revealed a marginally or significantly positive correlation between consistency and the rank order of the visual areas for both the Euclidean and non-Euclidean feature changes (*r* = .29*, t(5)* = 1.54, *p* = 0.092 for image stats, and *rs* > .33, *ts* > 2.34, *ps* < .044 for all other nonidentity features). Direct comparisons further confirmed that, for position, size and SF changes, the average consistency of V1 and V2 was lower than that of LOT and VOT (*ts* > 2.45, *ps* < .033). This difference was not significant for the image stats change (*t(5)* = 1.46, *p* = .10). In additional analyses, we showed that there was no eye position confound on consistency measures and that consistency in early visual areas was not a direct reflection of consistency at the image pixel level (see Supplementary Results).

Overall, these results show the existence of high consistency in object representational space across both Euclidean and non-Euclidean feature changes in higher visual areas. For the Euclidean feature changes, consistency was absent in lower visual areas, indicating a lack of representational uniformity across visual field in these areas. Nevertheless, over the course of processing, consistency gradually rose to become fairly high by the end of ventral processing. For the non-Euclidean feature changes, there was already a fair amount of consistency in lower visual areas, indicating some representational uniformity across these feature changes in these areas. While ventral visual processing further increased consistency across an SF change, the effect across image stats change was much weaker.

### The relationship among the three tolerance measures in human ventral visual areas

The separate analyses performed above reveals a largely similar response profile across rank-order preservation, cross-decoding success and representational consistency over the course of ventral visual processing. Across all three measures, there was a large increase from lower to higher visual areas for the two Euclidean feature changes; in comparison, the increase for the two non-Euclidean feature changes was relatively small. To quantify the coupling among these three measures, within each participant we correlated the values from the six brain regions between pairs of measures and then evaluated the results at the group level. For the two Euclidean feature changes, significant correlations were obtained for all pairs of correlations (*rs* > .51, *ts* > 2.46, *ps* < .024; one-tailed and corrected for the three pairwise correlations performed for each feature change; one-tailed t tests were used here to test for the difference in a specific direction; this applies to all subsequent t tests of the same type). For the two non-Euclidean feature changes, for the image stats change, all pairs of correlations were either marginally significant or significant (*r* = .34, *t(5)* = 2.01, *p* = .0504 for the correlation between cross-decoding and consistency; and *rs* > .50, *ts* > 3.82, *ps* < .0093, for the other two pairs of correlations). For the SF change, the correlation between rank-order and consistency was significant (*r* = .60, *t(10)* = 4.13, *p* = .0039), but those for the other two pairs were not (*rs* < .13, *ts* < .51, *ps* > .31).

Just as rank-order preservation at the single neuron level has been shown to be tightly correlated with cross-decoding at the neuronal population level across nonidentity feature changes (Li et al., 2009), here we showed for the first time that such as a correspondence is also present between these two measures in fMRI voxels. Moreover, for Euclidean feature changes, there was a very strong coupling among rank-order preservation, cross-decoding success, and consistency, with the values going up from lower to higher visual areas in a similar manner across these different measures. A similar, but weaker coupling across the three measures was also present for the non-Euclidean feature changes.

### Tolerance measures in CNNs

CNNs are considered as the current best models of the primate visual system due to their high object categorization performance and their ability to identify objects across a variety of identity preserving image transformations (Yamins & Dicarlo, 2016; Kriegeskorte, 2015). Does rank-order preservation exist in CNN units? What about representational consistency? What is the relationship among cross-decoding, rank-order preservation and consistency over the course of CNN visual processing? To answer these questions, we examined a total of 8 CNNs, including both shallower networks, such as Alexnet, VGG16 and VGG 19, and deeper networks, such as Densenet-201, Googlenet, Resnet-50 and Resnet-101 (Table 1). We also included a recurrent network, Cornet-S, that has been shown to capture the recurrent processing in macaque IT cortex with a shallower structure and has been argued to be the current best model of the primate ventral visual regions (Kubilius et al., 2019; Kar et al., 2019). All CNNs were pretrained with ImageNet images (Deng et al., 2009). Following previous studies (e.g., O’Connor & Chun, 2018; Taylor & Xu, 2021; Xu & Vaziri-Pashkam, 2021a & 2021b), we sampled from 6 to 11 mostly pooling layers of each CNN (see Table 1 for the specific CNN layers sampled). For each unit of each sampled CNN layer, we examined the averaged response from all the images of the same object category at each value of a given nonidentity feature, similar to how an fMRI category response was extracted (see Methods).

**Table 1.** CNNs and layers sampled.

| CNN name | Depth/Blocks | Layers | N of Layers Sampled | Sampled Layer Names and Locations (indicated in the parenthesis) |
| --- | --- | --- | --- | --- |
| Alexnet | 8 | 25 | 6 | 'pool1' (5), 'pool2' (9), 'pool5' (16), 'fc6' (17), 'fc7' (20), 'fc8' (23) |
| Cornet-S | 4 | 42 | 6 | 'V1_outpt' (8), 'V2_output' (18), 'V4_output' (28), 'IT_output' (38), 'decoder_avgpool' (39), 'decoder_output' (42) |
| Densenet-201 | 201 | 709 | 6 | 'pool1' (6), 'pool2_pool' (52), 'pool3_pool' (140), 'pool4_pool' (480), 'avg_pool' (706), 'fc1000' (707) |
| Googlenet | 22 | 144 | 6 | 'pool1-3x3_s2' (4), 'pool2-3x3_s2' (11), 'pool3-3x3_s2' (40), 'pool4-3x3_s2' (111), 'pool5-7x7_s1' (140), 'loss3-classifier' (142) |
| Resnet-18 | 18 | 72 | 6 | 'pool1' (6), 'res2b_relu' (20), 'res3b_relu' (36), 'res4b_relu' (52), 'pool5' (69), 'fc1000' (70) |
| Resnet-50 | 50 | 177 | 6 | 'max_pooling2d_1' (5), 'activation_10_relu' (37), 'activation_22_relu' (79), 'activation_40_relu' (141), 'avg_pool' (174), 'fc1000' (175) |
| Vgg-16 | 16 | 41 | 8 | 'pool1' (6), 'pool2' (11), 'pool3' (18), 'pool4' (25), 'pool5' (32), 'fc6' (33), 'fc7' (36), 'fc8' (39) |
| Vgg-19 | 19 | 47 | 8 | 'pool1' (6), 'pool2' (11), 'pool3' (20), 'pool4' (29), 'pool5' (38), 'fc6' (39), 'fc7' (42), 'fc8' (45) |

Despite differences in the exact architecture, depth, and presence/absence of recurrent processing, largely similar trajectories in rank-order preservation, cross-decoding, and consistency over the course of visual processing were observed for all 8 CNNs tested (Figure 4). For the two Euclidean feature changes, responses from the three measures were largely monotonic (i.e., either increasing or decreasing) over the course of visual processing. For position, both rank-order preservation and cross-decoding remained near zero for the first few layers sampled and then increased monotonically from mid to higher layers. Cross-decoding reached 1 by the end of CNN processing for every CNN tested, showing that CNNs are capable of fully generalizing their representations across a position change. Meanwhile, none of the rank-order preservation reached 1 by the end of processing. All 8 CNNs showed an overall high consistency, but with a downward trend from lower to higher layers, the opposite of what was seen in cross-decoding and rank-order preservation. For size, while cross-decoding increased steadily from lower to higher layers, rank-order preservation remained close to zero for the first few layers and then increased from mid to higher layers. The rise in rank-order preservation across layers thus lagged behind that of cross-decoding: even when rank-order preservation was zero or close to zero, significant cross-decoding was still obtained. While Densenet-201 and Resenet-18 were able to reach cross-decoding of 1 by the end of processing, the other CNNs failed to do so. As with the position change, all 8 CNNs showed a downward trend for consistency, going from 1 to dropping below .8 from lower to higher layers across all 8 CNNs (with a few dropping below .4). Similar CNN response patterns were obtained when the original, rather than the controlled, images were used for position and size changes (see Supplementary Figure 1), showing that the downward trend observed in both for consistency was not due to the controlled images used.

**Figure 4.**
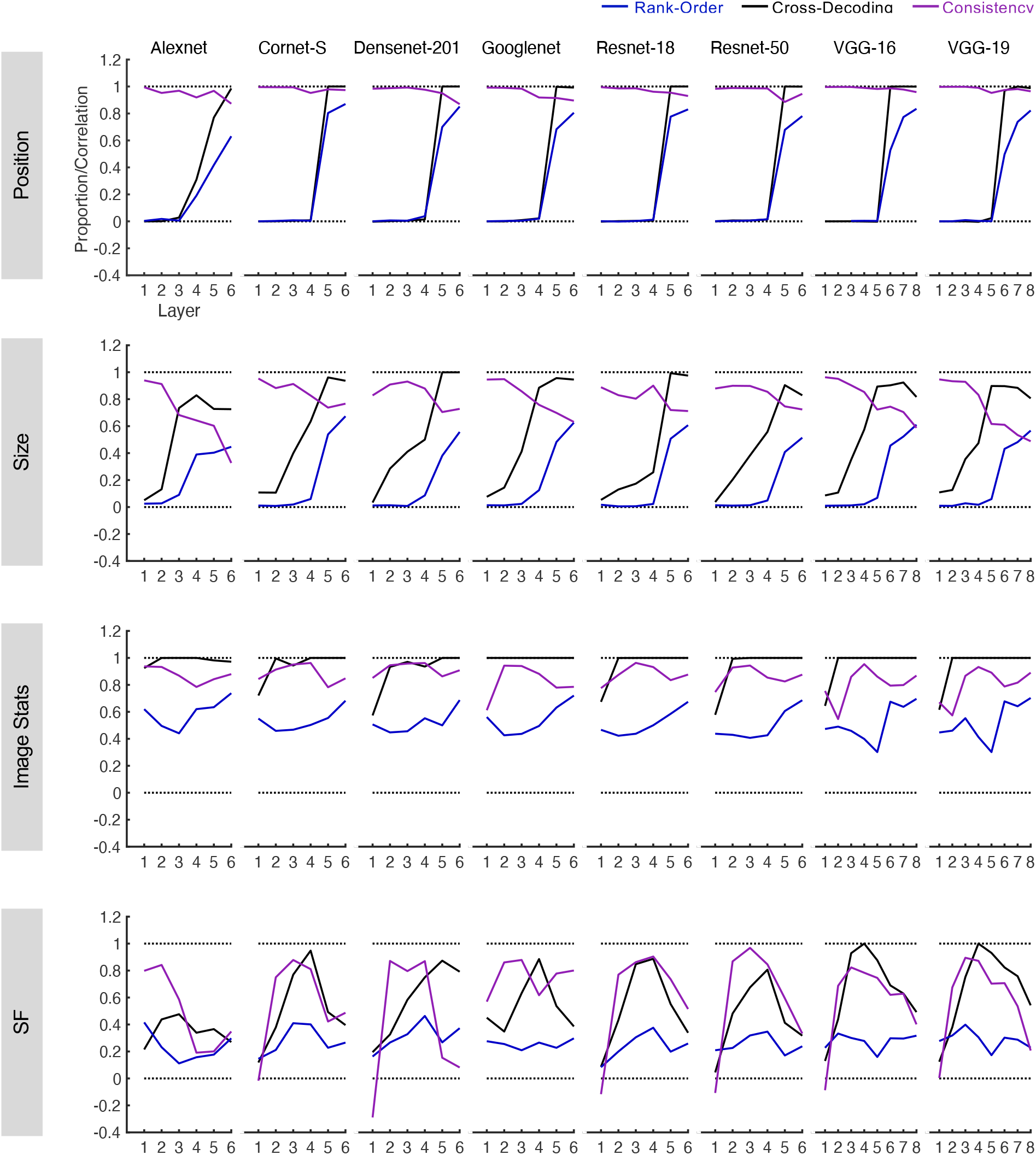
Rank-order preservation, cross-decoding success, and consistency from 8 CNNs pretrained for image classification across the two Euclidean (position and size) and the two non-Euclidean (image stats and SF) feature changes. These measures were generated in similar ways as the corresponding brain measures, except that correction for reliability was not applied as CNNs had no noise (see Methods).

Responses for the two non-Euclidean feature changes tended to show large fluctuations over the course of visual processing not seen in brain responses. For image stats change, for cross-decoding, except for Densenet-201 in which performance rose to 1 by the end of processing, for all others, performance rose quickly to 1 early on in processing. Although rank-order preservation also showed an overall increase from lower to higher layers, it was not monotonic, with a dip occurring in the middle layers that was absent in the cross-decoding results. Consistency tended to show an inverted U-shape across the layers, the opposite of what was seen with rank-order preservation. For SF change, none of the measures showed a rise from lower to higher layers, with both cross-decoding and consistency showing an inverted U-shape between lower and higher layers and with none of the CNN cross-decoding reaching close to 1 by the end of processing. These response patterns differed substantially from the largely monotonic increase in the corresponding brain responses.

Overall, these results show some significant decoupling among the three tolerance measures for each nonidentity feature in every CNN tested. This is different from the measures seen in human visual areas.

### Comparing tolerance measures between the brain and CNNs

To directly compare between the brain and CNN results, for each of the three tolerance measures, using Spearman rank correlation, we correlated the overall response profiles across visual hierarchy between the brain and CNNs. The resulting correlation coefficients are plotted against the upper and lower bounds of the noise ceiling of the brain data (Figure 5), with correlations below the lower bound indicating that a given CNN measure was significantly different from the corresponding measures in the human brain. For cross-decoding and rank-order preservation, measures from a number of CNNs reached the lower bound of the noise ceiling for human brain measures; however, for consistency, this was not the case. For the two Euclidean feature changes, no brain-CNN correlation for consistency reached the noise ceiling. For the two non-Euclidean feature changes, the vast majority of brain-CNNs correlations for consistency also failed to reach the noise ceiling. Overall, about 42% to 58% of the comparisons for a given CNN were significantly or marginally significantly below the noise ceiling (42% for Cornet, Densenet-201, Googlenet, and Resnet-18; 50% for VGG-19; and 58% for Alexnet, Resnet-50, and VGG -16; see Table 2 for the detailed stats; results were corrected for multiple comparisons for the three comparisons made within each CNN and each nonidentity feature change).

**Figure 5.**
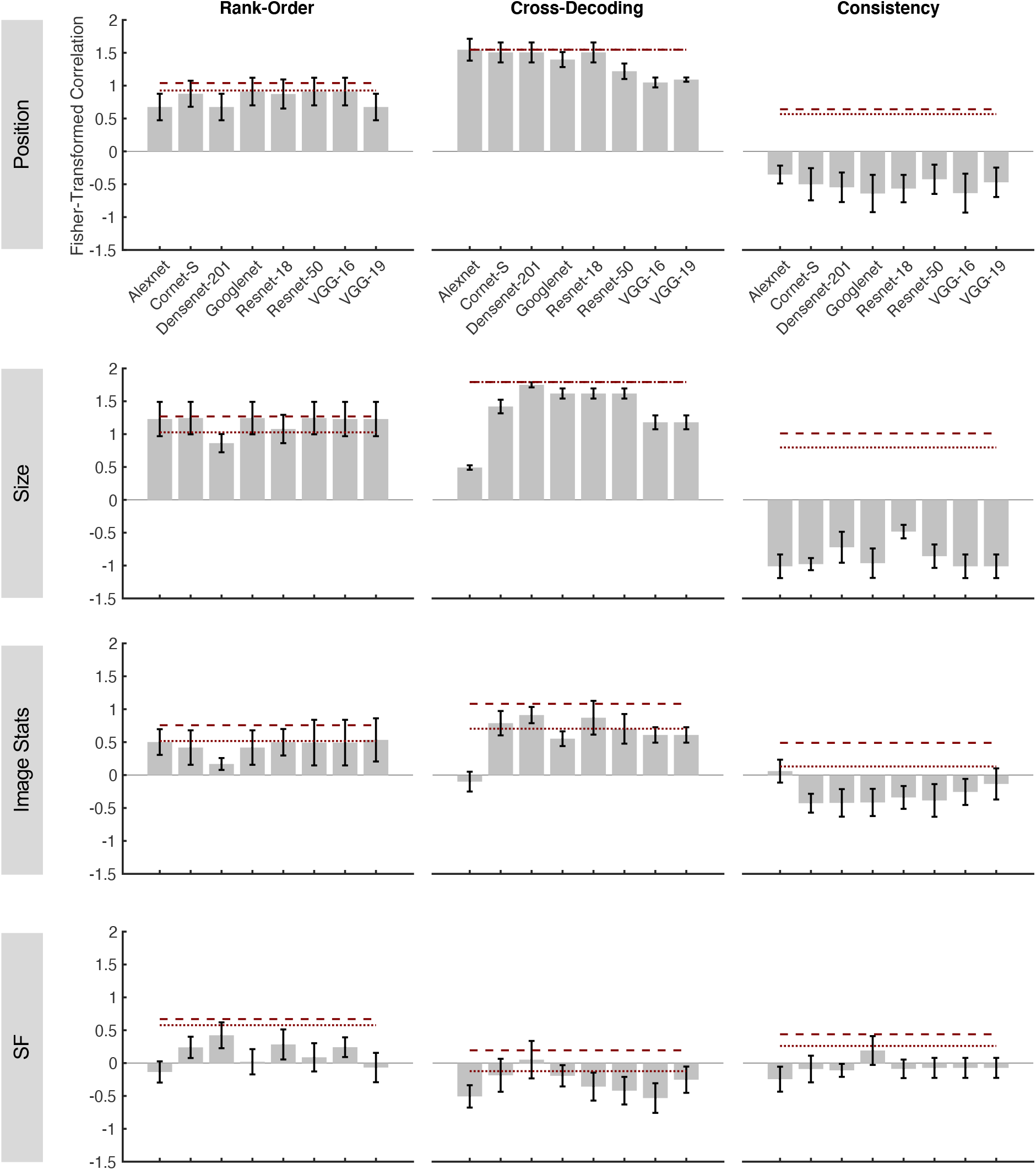
Brain and CNN correlations of response profiles for rank-order preservation, cross-decoding success and consistency across the two Euclidean (position and size) and the two non-Euclidean (image stats and SF) feature changes. The dashed and dotted dark red lines represent the upper and lower bounds of the noise ceiling, respectively, calculated from the brain data. To directly compare between the brain and CNN results, for each of the three measures, using Spearman rank correlation, we correlated the overall response profiles over the course of visual processing between a given CNN and the brain data from each human participant. The resulting averaged Fisher-transformed correlation coefficients were then plotted against the upper and lower bounds of the noise ceiling of the brain data, with correlations significantly below the lower bound indicating a given CNN measure being different from the corresponding brain measure.

**Table 2.** Brain and CNN correlations of response profiles for rank-order preservation (rank order), cross-decoding success (decoding), and consistency across the four types of nonidentity feature changes. To directly compare between the brain and CNN results, for each of the three measures, using Spearman rank correlation, the overall response profiles over the course of visual processing between a given CNN and each human participant were first correlated. The resulting Fisher-transformed correlation coefficients were then tested against the lower bound of the noise ceiling of the brain data using 1-tailed t tests, with correlations below the lower bound indicating a given CNN measure being significantly different from the corresponding measures from the human brain. Results were corrected for multiple comparisons for the three comparisons made within each CNN and each nonidentity feature change using the Benjamini-Hochberg method (Benjamini & Hochberg, 1995). t/p values are reported. Bolded texts indicates statistically significant results (*ps* < .05) and grey bolded texts indicates marginally significant results (.05 < *ps* < .1).

|  | <u>Position</u> |  |  | <u>Size</u> |  |  |
| --- | --- | --- | --- | --- | --- | --- |
|  | Rank Order | Decoding | Consistency | Rank Order | Decoding | Consistency |
| Alexnet | -1.25/.20 | <.001/.5 | **-6.77/<.001** | .77/.77 | **-38.48/<.001** | **-10.06/<.001** |
| Cornet-S | -.25/.41 | -.27/.60 | **-4.36/.0072** | .88/.79 | **-3.60/.0086** | **-19.27/<.001** |
| Densenet-201 | -1.25/.20 | -.27/.40 | **-4.95/.0039** | -1.18/.14 | -1.10/.24 | **-6.44/<.001** |
| Googlenet | -.076/.47 | -1.32/.18 | **-4.25/.0081** | .88/.79 | **-2.26/.048** | **-7.86/<.001** |
| Resnet-18 | -.24/.41 | -.27/.60 | **-5.44/<.001** | .24/.59 | **-2.26/.048** | **-12.48/<.001** |
| Resnet-50 | -.076/.47 | **-2.79/.023** | **-4.46/.0063** | .88/.79 | **-2.26/.048** | **-9.28/<.001** |
| Resnet-50-SIN | -.24/.62 | <.001/.5 | **-4.78/.0045** | -.57/.30 | **-2.26/.048** | **-13.14/<.001** |
| Resnet-50-SININ | -.24/.41 | -.27/.60 | **-5.87/<.001** | -.57/.30 | **-2.26/.048** | **-6.94/<.001** |
| Resnet-50-SININ-IN | -.076/.47 | -.27/.60 | **-4.83/.0045** | -.57/.30 | **-2.26/.048** | **-9.24/<.001** |
| VGG-16 | -.076/.47 | **-6.65/<.001** | **-4.06/.0049** | .77/.76 | **-5.80/<.001** | **-10.06/<.001** |
| VGG19 | -1.25/.13 | **-14.10/<.001** | **-4.63/.0027** | .77/.76 | **-5.80/<.001** | **-10.06/<.001** |
|  | <u>Image Stats</u> |  |  | <u>SF</u> |  |  |
|  | Rank Order | Decoding | Consistency | Rank Order | Decoding | Consistency |
| Alexnet | -.08/.47 | **-5.34/.0045** | -.41/.52 | **-4.41/<.001** | **-2.26/.025** | **-2.66/.020** |
| Cornet-S | -.38/.54 | .45/.67 | **-3.90/.018** | -2.08/.10 | -.25/.40 | **-1.73/.089** |
| Densenet-201 | **-3.86/.018** | 1.68/.92 | **-2.65/.033** | -.78/.35 | .61/.72 | **-3.74/.0069** |
| Googlenet | -.38/.36 | -1.36/.18 | **-2.64/.066** | **-2.90/.026** | -.44/.51 | -.32/.38 |
| Resnet-18 | -.09/.69 | .65/.73 | **-2.71/.063** | -1.28/.18 | -1.10/.15 | **-2.47/.054** |
| Resnet-50 | -.07/.71 | -.008/.50 | -2.08/.14 | **-2.27/.075** | **-1.41/.096** | **-2.19/.042** |
| Resnet-50-SIN | -.38/.54 | 1.43/.89 | **-2.65/.069** | -2.05/.11 | -.75/.24 | **-1.97/.060** |
| Resnet-50-SININ | -.38/.54 | 1.43/.89 | **-2.65/.069** | -.33/.37 | -.75/.24 | **-2.47/.054** |
| Resnet-50-SININ-IN | -.38/.54 | 1.43/.89 | **-3.25/.033** | **-3.31/.014** | **-1.41/.096** | **-2.47/.027** |
| VGG-16 | -.07/.47 | -.81/.35 | -1.95/.16 | **-2.24/.078** | **-1.81/.051** | **-2.19/.042** |
| VGG19 | .05/.52 | -.81/.35 | -1.12/.48 | **-2.89/.027** | -.65/.27 | **-2.19/.042** |

To examine the coupling among the three measures in CNNs and whether they showed the same coupling as those found in the brain, we compared the pairwise correlation coefficients of the three measures between the brain and CNNs (Figure 6). Specifically, we evaluated whether the pairwise correlations obtained from the CNNs were significantly outside the corresponding values obtained from the human brain (see Table 3 for the detailed stats). For the correlation between rank-order preservation and cross-decoding, a good number of CNNs showed a positive correlation no different from that of the brain, with VGG19 being the best (showing no difference for all four nonidentity feature changes) and Alexnet being the worse (showing significant differences for all four). For the correlation between rank-order preservation and consistency, no CNNs showed similar correlation as the brain for position, size and image stats changes. For the correlation between cross-decoding and consistency, no CNNs showed similar correlation with the brain for the two Euclidean feature changes. Overall, about 50% to 83% of the brain-CNN comparisons for a given CNN yield significant or marginally significant differences (50% for VGG-19; 58% for Densenet-201; 67% for Cornet-S, Resnet-18 and VGG-16; 75% for Googlenet, Resnet-50; and 83% for Alexnet; see Table 3 for the detailed stats; results were corrected for multiple comparisons for the three comparisons made within each CNN and each nonidentity feature change).

**Figure 6.**
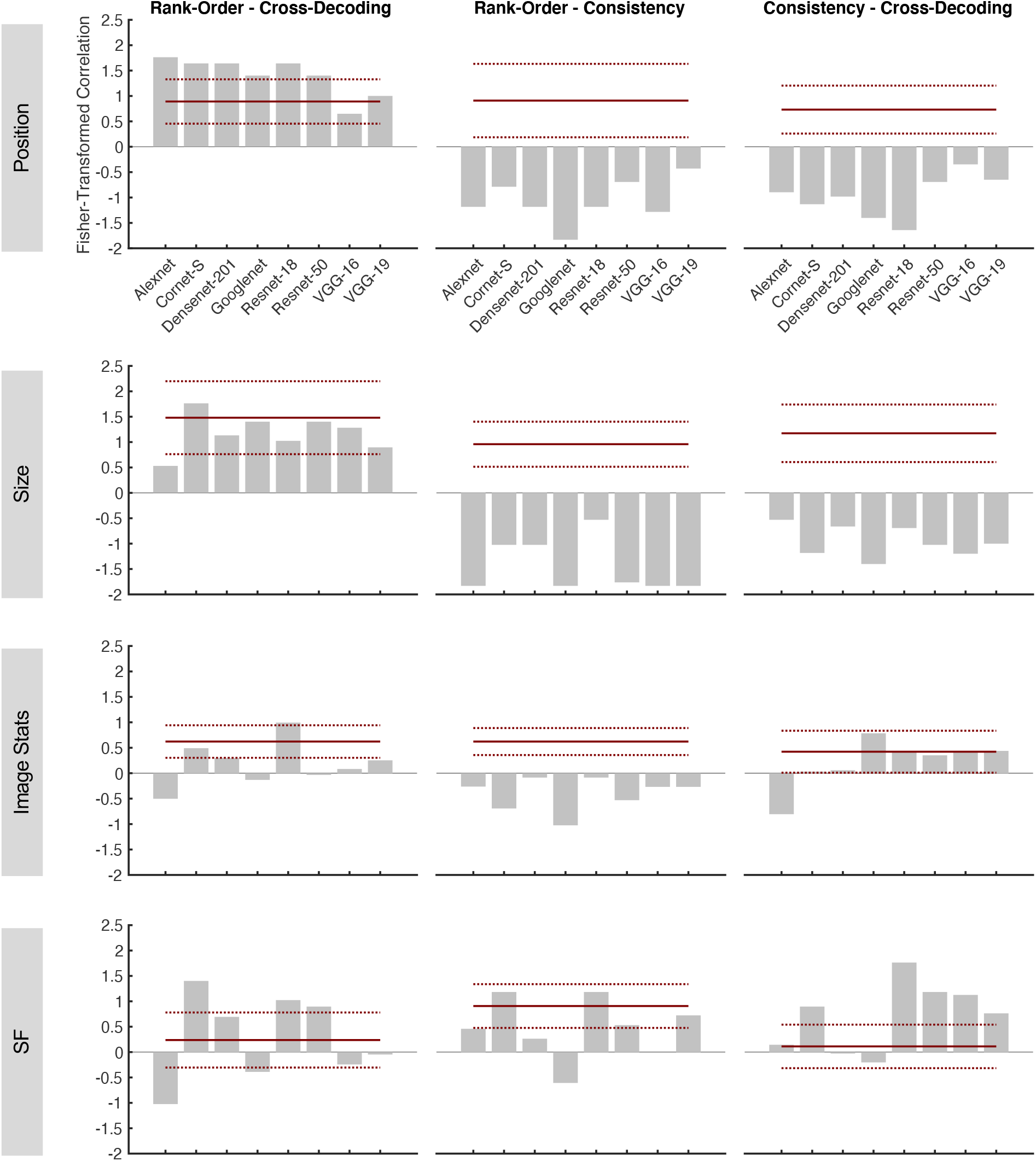
Brain and CNN comparisons of the coupling in response profiles among rank-order preservation, cross-decoding success, and consistency for the two Euclidean (position and size) and the two non-Euclidean (image stats and SF) feature changes. The solid dark red lines represent the mean correlation coefficients from the brain, with the dotted dark red lines indicating the 95% confidence intervals around the means. To compare the coupling in response profiles among these three measures between the CNNs and the brain, pairwise correlation coefficients of the response profiles from these three measures in the CNNs were plotted against the corresponding values from the brain. A CNN value significantly different from the corresponding brain value indicates a difference in coupling between the CNN and the brain. All values shown were Fisher z-transformed to ensure valid statistical comparisons.

**Table 3.** Brain and CNN comparisons of the coupling in response profiles among rank-order preservation (R), cross-decoding success (D), and consistency (C) for the four types of nonidentity feature changes. To examine whether the coupling in response profile among these three measures in CNNs differed from those from the brain, pairwise correlation coefficients of the response profiles from these three measures from each CNN were tested against the corresponding values from the brain using two-tailed t tests, with the CNN values significantly different from the brain values indicating a difference in coupling between the brain and CNN. Results were corrected for multiple comparisons for the three comparisons made within each CNN and each nonidentity feature change using the Benjamini-Hochberg method (Benjamini & Hochberg, 1995). Black bolded texts indicates statistically significant results (*ps* < .05) and grey bolded texts indicates marginally significant results (.05 < *ps* < .1). R-D, rank-order preservation and cross-decoding success; R-C, rank-order preservation and consistency; D-C, cross-decoding success and consistency.

|  | <u>Position</u> |  |  | <u>Size</u> |  |  |
| --- | --- | --- | --- | --- | --- | --- |
|  | R-D | R-C | D-C | R-D | R-C | D-C |
| Alexnet | **-3.92/.0078** | **5.67/.0039** | **5.67/.0019** | **2.59/.041** | **12.32/<.001** | **5.89/.0016** |
| Cornet-S | **-3.37/.045** | **4.60/.0056** | **4.60/.0037** | -.77/.47 | **8.75/<.001** | **8.15/<.001** |
| Densenet-201 | **-3.37/.015** | **5.67/.0039** | **5.67/.0019** | .95/.38 | **8.75/<.001** | **6.35/<.001** |
| Googlenet | **-2.29/.06** | **7.43/<.001** | **7.43/<.001** | .21/.84 | **12.32/<.001** | **8.90/<.001** |
| Resnet-18 | **-3.37/.015** | **5.67/.0039** | **5.67/.0019** | 1.25/.26 | **6.57/<.001** | **6.45/<.001** |
| Resnet-50 | **-2.29/.06** | **4.34/.0073** | **5.91/.0030** | .21/.84 | **12.01/<.001** | **7.60/<.001** |
| Resnet-50-SIN | **-3.92/.0078** | **5.24/.0029** | **8.85/<.001** | 1.59/.16 | **6.57/<.001** | **7.60/<.001** |
| Resnet-50-SININ | **-3.37/.015** | **4.11/.0095** | **6.60/<.001** | 1.59/.16 | **6.91/<.001** | **5.89/.0016** |
| Resnet-50-SININ-IN | **-4.22/.0055** | **5.24/.0029** | **7.74/<.001** | 1.59/.16 | **6.25/<.001** | **6.45/<.001** |
| VGG-16 | 1.09/.32 | **5.94/.0030** | **5.94/.0020** | .54/.61 | **12.32/<.001** | **8.20/<.001** |
| VGG19 | -.49/.64 | **3.63/.033** | **3.63/.017** | 1.59/.16 | **12.32/<.001** | **7.52/<.001** |
|  | <u>Image Stats</u> |  |  | <u>SF</u> |  |  |
|  | R-D | R-C | D-C | R-D | R-C | D-C |
| Alexnet | **6.89/.003** | **6.54/.0019** | **5.83/.0021** | **4.56/.0042** | 2.04/.11 | -.15/.89 |
| Cornet-S | .80/.46 | **9.72/p<.001** | 1.86/.18 | **-4.21/.0069** | -1.26/.24 | **-3.58/.0089** |
| Densenet-201 | 1.99/.15 | **5.23/.010** | 1.73/.14 | -1.65/.20 | **2.93/.051** | .64/.54 |
| Googlenet | **4.62/.0086** | **12.15/p<.001** | -1.71/.15 | **2.27/.074** | **6.89/p<.001** | 1.44/.18 |
| Resnet-18 | -2.27/.11 | **5.23/.010** | .025/.98 | **-2.84/.029** | -1.26/.24 | **-7.55/<.001** |
| Resnet-50 | **4.02/.020** | **8.51/p<.001** | .34/.75 | **-2.38/.062** | 1.71/.12 | **-4.90/<.001** |
| Resnet-50-SIN | -.81/.46 | **9.09/p<.001** | 1.46/.32 | .55/.90 | .052/.96 | **-2.66/.078** |
| Resnet-50-SININ | -.81/.46 | **9.09/p<.001** | 1.46/.32 | -.09/.93 | **2.35/.065** | **-4.17/.0072** |
| Resnet-50-SININ-IN | -.81/.46 | **9.72/p<.001** | 1.73/.21 | -1.34/.32 | 1.36/.21 | **-7.86/<.001** |
| VGG-16 | **3.31/.032** | **6.58/.0036** | -.075/.94 | 1.74/.12 | **4.13/.0039** | **-4.64/.0036** |
| VGG19 | 2.27/.11 | **6.58/.0036** | -.075/.94 | 1.03/.50 | .84/.42 | **-2.98/.048** |

Together, these results showed that there were similarities as well as large discrepancies between the brain and CNNs in the three tolerance measures. For the two Euclidean feature changes, both rank-order preservation and cross-decoding rose from around zero to around 1 over the course of processing. These characteristics and the coupling between these two measures resembled the measures from human visual areas. However, unlike the brain, consistency declined with processing, resulting in a decoupling between consistency and the other two measures as well as large discrepancies between CNN and brain measures. For the two non-Euclidean feature changes, CNNs showed large fluctuations in tolerance measures not seen in the brain and large deviations from the corresponding brain measures. CNNs thus do not appear to form brain-like transformation-tolerant visual object representations over the course of visual processing.

### Training a CNN with stylized object images

Although CNNs are believed to explicitly represent object shapes in the higher layers (Kriegeskorte, 2015; LeCun et al., 2015; Kubilius et al., 2016), emerging evidence suggests that CNNs may mostly use local texture patches to achieve successful object classification (Ballester & de Araújo, 2016, Gatys et al., 2017; Geirhos et al., 2019). However, when Resnet-50 is trained with stylized ImageNet images in which the original texture of every single image is replaced with the style of a randomly chosen painting, object classification performance significantly improves, relies more on shape than texture cues, and becomes more robust to noise and image distortions (Geirhos et al., 2019).

To test whether training with stylized object images would make CNNs exhibit more brain-like tolerance, we extracted responses from Resnet-50 pretrained with stylized ImageNet Images under three different training protocols (Geirhos et al., 2019). Comparing these responses with responses from Resnet-50 pretrained with the original ImageNet images, we found overall remarkedly similar results in all three tolerance measures (Supplementary Figure 2) as well as their correlations with the corresponding brain measures (Supplementary Figure 3; see Tables 2 and 3 for the detailed stats). The failure of Resnet-50 to exhibit more brain-like responses across nonidentity feature changes suggests that there are likely differences between the two that cannot be overcome by this type of training.

## Discussion

Forming transformation-tolerant object representations is essential for high-level primate vision. Here we provide a comprehensive documentation of three tolerance measures in human ventral visual cortex and 8 CNNs when objects undergo both Euclidean and non-Euclidean feature changes including position, size, image stats and SF.

### Rank-order preservation in fMRI voxels for objects across nonidentity feature changes in human ventral visual areas

We show here for the first time that robust response rank-order preservation exists in individual fMRI voxels in human higher visual areas for objects undergone both Euclidean and non-Euclidean feature changes. Such a finding has only been reported previously for isolated single neurons in macaque higher visual areas. This indicates the existence of functional smoothness at high-level vision such that neurons with a similar rank-order preservation profile are congregated together. To our knowledge, such an organization at the fMRI voxel meso-scale level has not been reported before. With fMRI, we additionally document for the first time the emergence of rank-order preservation over the ventral visual areas, as previous neurophysiological studies only examined very few areas due to challenges associated with recording from multiple areas (e.g., Li et al., 2009 recorded from IT but simulated results from V1 and Rust et al. 2010 only recorded from V4 and IT). Our examination of both Euclidean and non-Euclidean features further revealed both differences as well as similarities in rank-order preservation to these feature changes. At lower visual areas, while rank-order preservation was low for the two Euclidean feature changes, it was already fairly high for the two non-Euclidean feature changes. Thus, it is not the case that rank-order preservation is a unique property of high-level vision. But rather, rank order could be maintained to some extent in lower visual areas for certain nonidentity feature changes. That being said, rank-order preservation increased from lower to higher visual areas for both the Euclidean and non-Euclidean features examined, showing that over the course of ventral processing, object response rank order become more aligned across a nonidentity feature change.

As in a previous modeling and simulation study with single neurons (Li et al., 2009), we also observed a close correspondence between rank-order preservation at the individual voxel level and cross-decoding success at the voxel population level for position, size and SF changes over the course of visual processing, with both increasing from lower to higher visual areas, respectively. We did find one instance (i.e., in image stats change) where this correspondence did not hold. This could be due to decoding being an overall less sensitive measure of the representational space, with different representational structures capable of producing similar decoding results so long as the representations are on the correct side of the decision boundary.

### Representational consistency for objects across nonidentity feature changes in human ventral visual areas

In higher visual areas, both object identity and nonidentity information can be independently read out (e.g., Hung et al., 2005; Schwarzlose et al., 2008; Sayres & Grill-Spector, 2008; Carlson et al., 2011; Cichy et al., 2011; Hong et al., 2016; Vaziri-Pashkam et al., 2019; Mocz et al., 2021). This requires orthogonal feature representations and object representational consistency in the feature space, with the similarity among a set of objects well preserved across a non-identity feature change. Here we report for the first time that high consistency indeed exists in higher ventral visual areas for both the Euclidean and non-Euclidean feature changes tested here, showing that the object representational space is well aligned across feature changes to support independent representations of object identity and nonidentity information towards the end of ventral processing. For the two Euclidean feature changes involving position and size, despite there being high consistency at the image input level (due to the uniformity of the image space, see Supplementary Results), consistency was absent in lower visual areas, indicating a lack of representational uniformity across visual field in these areas. Nevertheless, over the course of processing, consistency gradually rose to become fairly high by the end of ventral processing. For the non-Euclidean feature changes, there was already a fair amount of consistency in lower visual areas, indicating some representational uniformity across these feature changes in these areas. While ventral visual processing further increased consistency across an SF change, the effect across image stats change was weaker. Overall, consistency showed a close coupling with rank order across all feature changes and with cross-decoding for three of the four feature changes; the coupling between consistency and cross-decoding in SF was not significant, likely due to the same reason as discussed above.

It may come as a surprise to some that there is a lack of uniformity in feature representation across space in early visual areas. Because neurophysiological studies can only sample a limited number of neurons, uniformity in early visual areas has not been directly tested. The existence of hyper-columns in V1 does not require the distribution of features to be uniform across space. In fact, it is easier to have non-uniformity than uniformity during development. Prior studies have identified a host of systematic non-uniformities in representation in early visual areas, such as the cortical magnification factor (Daniel & Whitteridge, 1961), finer spatial and greater curvature representation in fovea than periphery (Srihasam, et al., 2014), radial bias in orientation representation (Sasaki, et al, 2006), and spatial field bias (Silson et al, 2018). Some of these factors could contribute to the non-uniformity in feature representation in early visual areas. We show here that, over the course of ventral processing, such non-uniformity gradually disappears. Achieving tolerance in visual processing thus can also be viewed as removing non-uniformity in feature representation across different values of nonidentity features.

### Comparing tolerance measures between the brain and CNNs

We observed similarities as well as large discrepancies between the brain and CNNs in the three tolerance measures. For the two Euclidean feature changes, both rank-order preservation and cross-decoding in CNNs rose from around zero to around 1 over the course of processing. These characteristics and the coupling between these two measures resembled the measures from human visual areas. However, unlike the brain, CNNs showed a downward trend for consistency. This downward trend was much more prominent for the size than position change, going from 1 to dropping below .8 from lower to higher layers (with a few CNNs dropping below .4). In comparison, consistency rose from around .35 to around .95 from lower to higher human visual areas. This resulted in a decoupling between consistency and the other two tolerance measures in CNNs as well as large discrepancies between CNN and brain measures. For the two non-Euclidean feature changes, CNNs showed large fluctuations in tolerance measures not seen in brain responses, decoupling among the three tolerance measures, and large deviations from the corresponding brain measures.

We observed similar performance among the 8 CNNs tested despite differences in the exact architecture, depth, and presence/absence of recurrent processing. For example, the recurrent CNN we tested, Cornet-S, which was designed to closely model ventral visual processing, did not outperform the other CNNs. Furthermore, training a CNN with stylized images did not improve performance either. Overall, across the 8 CNNs, about 42% to 83% of the brain-CNN tolerance comparisons showed significant or marginally significant differences.

Lower layers of CNNs are designed to have uniformity across space. This and the heavy use of convolution to capture translational invariance (LeCun, 1989) could give rise to consistency in lower layers across position and size changes even when rank-order preservation and cross-decoding success are low (i.e., due to the small receptive field size, a unit may only respond to objects shown at one position and thus could not maintain rank-order and cross-decode objects across positions). While visual processing in human ventral regions resulted in a better alignment of objects in the representational space across nonidentity feature changes, CNNs appeared to lose some of the consistency they were given over the course of processing. Perhaps we can introduce nonuniformity across space in CNN lower layers to lower consistency across position and size changes to match that seen in human lower visual areas. However, doing so provides no guarantee that consistency would rise instead of dropping further over the course of processing. The core issue here is that visual processing in CNNs fails to increase or maintain consistency across nonidentity feature changes. This shows that despite CNNs’ success in classifying objects under different viewing conditions, they do not form brain-like transformation-tolerant object representations at the final stages of visual processing.

### Implications for CNNs

Our results add to the list of differences that have been reported between the brain and CNNs, such as CNNs’ ability to capture lower, but not higher, levels of visual representational structures of real-world objects and their inability to represent artificial objects like the human brain (Xu & Vaziri-Pashkam, 2021a), differences in the representational strength of non-identity features across visual processing (Xu & Vaziri-Pashkam, 2021b), their ability to explain only about 60% of the response variance of macaque V4 and IT (Cadieu et al., 2014; Yamins et al., 2014; Kar et al. 2019; Bashivan et al., 2019; Bao et al., 2020), their usage of different features compared to the primate brain for object recognition (Ballester & de Araujo, 2016, Ulman et al., 2016; Gatys et al., 2017; Baker et al., 2018; Geirhos et al., 2019; Jacob et al., 2021), and their susceptibility to the negative impact of adversarial images (Serre, 2019). Our findings are consistent with another study showing that while visual representations in humans exhibit tolerance to image changes across time, those from CNNs do not (Henaff et al., 2019).

While some of these discrepancies may be superficial (Firestone, 2020), our finding that the development of tolerance differs between the brain and CNNs and that consistency is not preserved in CNNs across feature changes at higher levels of visual processing indicates that CNNs use a fundamentally different computational algorithm to achieve tolerance at higher levels of visual representation. With its vast computing power, CNNs likely associate different instances of an object via a brute force approach (i.e., by simply remembering and grouping all instances of an object encountered under the same object label) without increasingly preserving the relationships among the objects across nonidentity feature changes to form brain-like tolerance. Indeed, CNNs can achieve a near perfect classification accuracy even when image labels were randomly shuffled (Zhang et al. 2016), demonstrating their ability to memorize associations between images and random class labels. While this is one way to achieve tolerance, it requires a large number of training data and has a limited ability to generalize to objects not included in training, the two major drawbacks associated with the current CNNs (Serre, 2019).

The formation of transformation-tolerant object representations in the primate brain has been argued to be critical in facilitating information processing and learning by reducing the number of training examples needed while at the same time increasing the generalizability from the trained images to new instances of an object and a category (Tacchetti et al., 2018). Even if CNNs were to use a fundamentally different, but equally viable, computational algorithm to solve the tolerance problem compared to the primate brain, implementing a brain-like algorithm may nevertheless help them overcome the two major drawbacks mentioned above. That being said, making CNNs more brain like has its own practical advantages: as long as CNNs “see” the world differently from the human brain, they will make mistakes that are against human prediction and intuition. Allowing CNNs to capture the nature of human vision and then improve upon it will not only ensure the reliability of the devices powered by CNNs, such as in self-driving cars, but also, ultimately, our trust in using such an information processing system. Thus, in addition to benchmarking object recognition performance, it may be beneficial for future CNN architectures and/or training regimes to explicitly improve object representational consistency across nonidentity feature changes at higher levels of CNN visual processing. Doing so may push forward the next leap in model development and make CNNs not only better models for object recognition but also better models of the primate visual brain.

## Materials and Methods

### fMRI Experimental Details

Details of the fMRI experiments have been described in previously published studies (Vaziri-Pashkam & Xu, 2019; Vaziri-Pashkam et al., 2019). They are summarized here for the readers’ convenience.

Seven (four females), seven (four females), six (four females) and ten (five females) healthy human participants with normal or corrected to normal visual acuity, all right-handed, and aged between 18-35 took part in the position, size, image stats and SF experiments, respectively. All participants gave their informed consent prior to the experiment and received payment for their participation. The experiment was approved by the Committee on the Use of Human Subjects at Harvard University. Each experiment was performed in a separate session lasting between 1.5 and 2 hours. Each participant also completed two additional sessions for topographic mapping and functional localizers. MRI data were collected using a Siemens MAGNETOM Trio, A Tim System 3T scanner, with a 32-channel receiver array head coil. For all the fMRI scans, a T2*-weighted gradient echo pulse sequence with TR of 2 sec and voxel size of 3 mm x 3 mm x 3 mm was used. FMRI data were analyzed using FreeSurfer (surfer.nmr.mgh.harvard.edu), FsFast (Dale et al., 1999) and in-house MATLAB codes. FMRI data preprocessing included 3D motion correction, slice timing correction and linear and quadratic trend removal. Following standard practice, a general linear model was then applied to the fMRI data to extract beta weights as response estimates.

In the position experiment, we tested position tolerance and presented images either above or below the fixation (Figure 2b). We used cut-out grey-scaled images from eight real-world object categories (faces, bodies, houses, cats, elephants, cars, chairs, and scissors) and modified them to occupy roughly the same area on the screen (Figure 2a). For each object category, we selected ten exemplar images that varied in identity, pose and viewing angle to minimize the low-level similarities among them. Participants fixated at a central red dot throughout the experiment. Eye-movements were monitored in all the fMRI experiments to ensure proper fixation. During the experiment, blocks of images were shown. Each block contained a random sequential presentation of ten exemplars from the same object category shown either all above or all below the fixation. To equate low-level image differences among the different categories, controlled images were shown. Controlled images were generated by equalizing contrast, luminance and spatial frequency of the images across all the categories using the shine toolbox (Willenbockel et al., 2010, see Figure 2b). All images subtended 2.9° x 2.9° and were shown at 1.56° above the fixation in half of the 16 blocks and the same distance below the fixation in the other half of the blocks. Each image was presented for 200 msec followed by a 600 msec blank interval between the images. Participants detected a one-back repetition of the exact same image. This task engaged participants’ attention on the object shapes and ensured robust fMRI responses. Two image repetitions occurred randomly in each image block. Each experimental run contained 16 blocks, one for each of the 8 categories in each of the two image positions. The order of the eight object categories and the two positions were counterbalanced across runs and participants. Each block lasted 8 secs and followed by an 8-sec fixation period. There was an additional 8-sec fixation period at the beginning of the run. Each participant completed one scan session with 16 runs for this experiment, each lasting 4 mins 24 secs.

In the size experiment, we tested size tolerance and presented images either in a large size (5.77° x 5.77°) or small size (2.31° x 2.31°) centered at fixation (Figure 2b). As in the position experiment, controlled images were used here. Half of the 16 blocks contained small images and the other half, large images. Other details of the experiment were identical to that of the position experiment.

In the image stats experiment, we tested image stats tolerance and presented images at fixation either in the original unaltered format or in the controlled format (subtended 4.6° x 4.6°) (Figure 2b). Half of the 16 blocks contained original images and the other half, controlled images. As described earlier, the controlled images were generated by equalizing contrast, luminance and spatial frequency of the images across all the categories using the shine toolbox (Willenbockel et al., 2010, see Figure 2b). Other details of the experiment were identical to that of position experiment.

In the SF experiment, only six of the original eight object categories were included and they were faces, bodies, houses, elephants, cars, and chairs. Images were shown in 3 conditions: Full-SF, High-SF, and Low-SF (Figure 2b). In the Full-SF condition, the full spectrum images were shown without modification of the SF content. In the High-SF condition, images were high-pass filtered using an FIR filter with a cutoff frequency of 4.40 cycles per degree. In the Low-SF condition, the images were low-pass filtered using an FIR filter with a cutoff frequency of 0.62 cycles per degree. The DC component was restored after filtering so that the image backgrounds were equal in luminance. Each run contained 18 blocks, one for each of the category and SF condition combination. Each participant completed a single scan session containing 18 experimental runs, each lasting 5 minutes. Other details of the experiment design were identical to that the position experiment. Only the results from the High-SF, and Low-SF conditions were included in the present analysis.

We examined responses from independently localized early visual areas V1 to V4 and higher visual processing regions LOT and VOT (Figure 2c). V1 to V4 were mapped with flashing checkerboards using standard techniques (Sereno et al., 1995). Following the detailed procedures described in Swisher et al. (2007) and by examining phase reversals in the polar angle maps, we identified areas V1 to V4 in the occipital cortex of each participant (see also Bettencourt & Xu, 2016) (Figure 2c). To identify LOT and VOT, following Kourtzi and Kanwisher (2000), participants viewed blocks of intact object and scrambled object images. These two regions were then defined as a cluster of continuous voxels in the lateral and ventral occipital cortex, respectively, that responded more to the intact than to the scrambled object images. LOT and VOT loosely correspond to the location of LO and pFs (Malach et al., 1995; Grill-Spector et al.,1998; Kourtzi & Kanwisher, 2000) but extend further into the temporal cortex in an effort to include as many object-selective voxels as possible in occipito-temporal regions.

To generate the fMRI response pattern for each ROI in a given run, we first convolved an 8-second stimulus presentation boxcar (corresponding to the length of each image block) with a hemodynamic response function to each condition; we then conducted a general linear model analysis to extract the beta weight for each condition in each voxel of that ROI. These voxel beta weights were used as the fMRI response pattern for that condition in that run. Following Tarhan and Konkle (2020), we selected the top 75 most reliable voxels in each ROI for further analyses (by including voxels showing consistent responses for all the stimulus conditions across the odd and even halves of the data). This was done by splitting the data into odd and even halves, averaging the data across the runs within each half, correlating the beta weights from all the conditions between the two halves for each voxel, and then selecting the top 75 voxels showing the highest correlation. We used the same fMRI analysis procedure in a prior study and found that CNNs could account for 100% of the representational variance in early visual areas and 60% in higher visual areas (Xu & Vaziri-Pashkam, 2021a). The latter result is on par with results obtained from neurophysiology studies.

### CNN Details

We included 8 CNNs in our analyses (see Table 1). They included both shallower networks, such as Alexnet, VGG16 and VGG 19, and deeper networks, such as Densenet-201, Googlenet and Resnet-50. We also included a recurrent network, Cornet-S, that has been shown to capture the recurrent processing in macaque IT cortex with a shallower structure (Kubilius et al., 2019; Kar et al., 2019). This CNN has been recently argued to be the current best model of the primate ventral visual processing regions (Kar et al., 2019). All the CNNs used were trained with ImageNet images (Deng et al., 2009). Note that to streamline the analyses, we only included 8 of the 14 CNNs previously examined in our brain-CNN comparison analyses (Xu & Vaziri-Pashkam, 2021a & 2021b).

To understand how the specific training images would impact CNN representations, besides CNNs trained with ImageNet images, we also examined Resnet-50 trained with stylized ImageNet images (Geirhos et al., 2019). We examined the representations formed in Resnet-50 pretrained with three different procedures (Geirhos et al., 2019): trained only with the stylized ImageNet Images (RN50-SIN), trained with both the original and the stylized ImageNet Images (RN50-SININ), and trained with both sets of images and then fine-tuned with the stylized ImageNet images (RN50-SININ-IN).

Following O’Connor and Chun (2018), we sampled between 6 and 11 mostly pooling and FC layers of each CNN (see Table 1 for the specific CNN layers sampled). Pooling layers were selected because they typically mark the end of processing for a block of layers before information is pooled and passed on to the next block of layers. When there were no obvious pooling layers present, the last layer of a block was chosen. It has been shown previously that such a sampling procedure captures the evolution of the representation trajectory fairly well, if not fully, as adjacent layers exhibit identical or very similar representations (Taylor & Xu, 2021).

Cornet-S and the different versions of Resnet-50 were implemented in Python. All other CNNs were implemented in Matlab. No averaging of space and feature was done for the CNN output. Output from all CNNs were analyzed and compared with brain responses using Matlab.

### Data Analyses

#### Single fMRI voxel response rank-order preservation across nonidentity feature changes

In previous neurophysiological studies, rank-order correlation was never directly computed due to the limited number of objects tested (e.g., three in Li et al., 2009). But rather, rank-order preservation was assessed via a separability index which computes the correlation between the actual neuronal responses and its predicted responses across nonidentity feature changes assuming orthogonal representations of the identity and the nonidentity features (Brincat & Connor 2004; Janssen et al. 2008; Li et al., 2009). This index was inherited from earlier research studying the independent tuning of different features in sensory areas (e.g., the tuning dynamics of spatial frequency and orientation in area V1 in Mazer et al., 2002). The index is therefore not specifically developed to capture rank-order preservation across feature changes in high-level vision. Rank-order preservation, by definition, only requires the rank order in response amplitudes for the different objects to be maintained after a nonidentity feature change. It does not imply a strictly linear relationship as required by an orthogonal representation. In fact, Li et al. (2009) showed through simulation that whether or not orthogonality is maintained does not affect performance on invariant recognition tasks. With the separability index, it is also unclear how we may correct for differences in measurement noises across the different brain areas to compare the indices across brain areas.

In our study, because we included either 6 categories (for the SF change) or 8 categories (for the other three nonidentity feature change), we have sufficient number of categories to allow us to directly compute the Spearman rank-order correlation across a given feature change. Moreover, by using a split-half approach, we were able to correct for differences in measurement noise across the different brain areas to allow us to make valid comparisons across brain areas. Specifically, we divided the data into odd and even halves and averaged the responses within each half for each voxel and each stimulus condition (see also Xu & Vaziri-Pashkam, 2021a). For each of the 75 most reliable voxels in an ROI, we then correlated its response to all the object categories across the two values of a nonidentity feature and across the two halves of the data using Spearman rank correlation and took the average across both directions of the feature change as the raw rank order correlation for that voxel (e.g., correlating the category responses for odd run upper position with those for even run lower position and vice versa, and then taking the average of these two correlations). We averaged these raw rank order correlations across all the selected voxels in a given ROI for each participant. To calculate the reliability of the rank correlation for each ROI, for each voxel, we correlated the category responses within the same value of a nonidentity feature across the two halves of the data using Spearman rank correlation and took the average of the correlations obtained from both values of a given feature as the reliability measure for that voxel (e.g., correlating the category responses for odd run upper position with those for even run upper position and correlating those for odd run lower position with those for even run lower position, and then taking the average of these two correlations). We then averaged these reliability measures across all selected voxels in a given ROI and across all participants to derive an ROI-specific reliability measure. The final corrected rank order correlation was computed as the raw rank order correlation from each participant for that ROI divided by the group averaged reliability measure for that ROI. This was done separately for all ROIs examined and for each participant. The results were then evaluated at the group level using statistical tests.

#### Decoding object categories within and across nonidentity feature changes in the human ventral visual areas

To examine if object category information present at one value of a nonidentity feature can be generalized to another value, and how this effect may emerge from human lower to higher ventral visual areas, in this analysis, we measured category decoding accuracy with no feature change (within-decoding; e.g., both training and testing were done for objects shown at the same position) and across a feature change (cross-decoding; e.g., training was done for objects at one position and testing was done for objects at another position). Following Kamitani and Tong (2005) and our previous studies (Vaziri-Pashkam & Xu, 2017 & 2019; Vaziri-Pashkam et al., 2019), for the 75 most reliable voxels within each brain region, we first applied z-normalization to the averaged pattern for each condition in each ROI in each run to remove amplitude differences between conditions, ROIs, and runs. From the n-1 runs of the data, we trained a support vector machine (SVM) classifier using LIBSVM (Chang & Lin, 2011) to discriminate fMRI response patterns from a pairs of object categories at one value of a nonidentity feature. The trained classifier was then asked to discriminate the same category pair in the left-out data from the nth run both with no feature change (within-decoding) and across a feature change (cross-decoding). The left-out data was rotated with an n-fold cross-validation procedure. This analysis was performed for each pair of object categories and across both values of a nonidentity feature. The results were then averaged within each ROI for each participant.

To examine how decoding generalization changes over the course of visual processing, we also computed a decoding proportion score by first subtracting 0.5 from the within- and cross-decoding accuracies and then taking the ratio of the two resulting values. A proportion score of 1 indicates equally good decoding performance within and across a feature change and a complete generalization of category representation across the feature change; a proportion score of 0 on the other hand indicates a complete failure to generalize across the feature change.

#### Representational consistency for object categories across nonidentity feature changes in the human ventral visual areas

As with the rank order preservation measures, to account for differences in measurement noise across the different ROIs to allow us to make valid comparisons across ROIs, using the same split-half approach as above, for the 75 most reliable voxels in each ROI of each run, we averaged the fMRI response patterns within the odd and within the even runs, and then applied z-normalization to the averaged pattern for each condition in each ROI in each half of the runs to remove amplitude differences between conditions and ROIs.

To determine the extent to which object category representations were consistent across a nonidentity feature change in an ROI, for each participant, within each half of the data, within each value of a feature change, we computed all pairwise Euclidean distances from the averaged z-normalized patterns of the object categories to generate a representational dissimilarity matrix (RDM). We then correlated these RDMs across the two values of a nonidentity feature across the two halves of the data using Spearman rank correlation and took the average across both directions of the change as the raw RDM correlation (e.g., correlating odd run upper position with even run lower position and vice versa, and then taking the average of these two correlations). We calculated the reliability of RDM correlation by correlating the RDMs within the same value of a nonidentity feature across the two halves of the data using Spearman rank correlation and took the average across the two values of the nonidentity feature as the reliability measure (e.g., correlating odd run upper position with even run upper position and correlating odd run lower position with even run lower position, and then taking the average of these two correlations). The reliability measures from individual participants were averaged to generate a group-level ROI-specific reliability measure. The final corrected RDM correlation for each participant was computed as the raw RDM correlation for a given ROI divided by the group averaged reliability measure for that ROI. This was done separately for all the ROIs and for each participant. These corrected RDM correlations were our consistency measures.

In several participants the raw RDM correlation was greater than the absolute value of the reliability measure, yielding the corrected RDM correlation to be outside the range of [-1, 1]. Since correlation should not exceed the range of [-1, 1], any values exceeding the range were replaced by the closest boundary value (1 or -1). Without such a correction we obtained very similar line plots as those shown in Figure 3, but with a few large error bars due to a few excessively large values (greater than 10) obtained during the RDM normalization process. This mainly occurred for the early visual areas for the position change, likely due to the small size of the visual stimuli used and consequently nosier neural measures obtained. The actual values would likely be lower than 1 (based on the mean of the other participants). To ensure that we did not introduce bias in the above analysis procedure, in a further analysis, instead of computing a normalized value for each participant, to deal with measurement noise more effectively, we averaged all values across participants first before computing a normalized value. Specifically, for each ROI, we averaged the raw RDM correlations across participants to compute a group averaged raw RDM correlation. This correlation was then normalized by dividing its value by the corresponding group average reliability score. To derive error bars to better estimate the variance of these measures, instead of averaging across all the participants for a feature change, we also averaged across *n-1* participants at a time and then rotated the left-out participant to obtain *n* separate measures. From these *n* measures, we calculated the mean and variance. These results are shown in Supplementary Figure 4. They replicated our individual participant normalized results, with low values in earlier regions and high values in higher regions. All results were also similar when no reliability correction was applied to the data (Supplementary Figure 4).

#### CNN rank order preservation, cross-decoding success, and representational consistency analyses

For each unit of a given CNN layer, we extracted the response of each object image in each category at each value of a nonidentity feature change. We then averaged the output of all the images from the same category at the same value of the nonidentity feature to be the unit response for that category at that feature value. In all the analyses, we only included units that participated in category representation by showing differential responses to the different object categories for at least one value of a nonidentity feature (i.e., the standard deviation across all the categories for at least one value of the nonidentity feature was nonzero). This was necessary in order for the rank-order preservation analysis to be valid. A large number of the units at the lower layers would only respond to the background and not to the objects in the images. As such, including these units and (after adding a small amount of noise to each unit to avoid nondefinable correlations) would result in a large number of zero rank-order correlations. However, these zero rank-order correlations would not reflect a lack of rank-order preservation across a nonidentity feature change, but rather due to the trivial reason of these units not being involved in object representation. Consequently, including the results of these units in computing the average rank-order correlation would significantly distort the results. Although the inclusion of these nonresponsive units would not distort cross-decoding accuracy and consistency measures (since the decision boundary for decoding is drawn at the most informative direction and nonresponsive unit dimensions would not affect consistency calculation), to be consistent across all three tolerance measures, the same set of responsive units as determined above were included in all three measures.

To perform the rank order analysis, because a given unit could exhibit identical responses to all the categories at one value of a nonidentity feature (e.g., if the unit was only selective to images shown at the top position, then its responses to images shown at the bottom position would be identical for all the categories), to avoid nondefinable correlations, we added a very small amount of random noise to a unit’s response to each category, performed the rank order correlation as we did with the brain data, repeated this procedure 100 times, and averaged the resulting correlation coefficients as the final rank order correlation value for that unit. The rest of the rank-order analysis steps were identical to those used for the brain data.

To perform the category decoding analysis within and between the two values of a nonidentity feature, we constructed CNN responses to simulate the brain data. Specifically, we collected CNN responses for the same number of “runs” as the corresponding fMRI experiment (18 runs for the SF and 12 runs for all the other three nonidentity features). Each “run” contained the category responses averaged over 10 exemplars for each category from each value of a nonidentity feature. Following the design of the fMRI experiments, the 10 exemplars in each “run” contained two exemplar repetitions. This enabled responses from the different “runs” to be slightly different. Due to a lack of noise, within-decoding was 1 for all the layers in all the CNNs tested. We thus directly reported the results of cross-decoding without first normalizing it by within-decoding accuracy. The rest of the decoding analysis steps were identical to those used for the brain data.

For the consistency analysis, we z-normalized the unit responses to generate the CNN layer response pattern for a given object category at each value of the nonidentity feature, similar to how fMRI category responses were extracted. As with the fMRI responses, Euclidean distance between the response patterns of a pair of categories sharing the same value of the nonidentity feature was then calculated to generate a category RDM for each value of a nonidentity feature. As in the rank-order preservation analysis, to avoid nondefinable correlations, we added a very small amount of random noise to each cell of the RDM, performed the RDM correlation across the two values of a nonidentity feature, repeated this procedure 100 times, and averaged the resulting correlation coefficients as the final consistency measure for a given CNN layer. The rest of the consistency analysis steps were identical to those used for the brain data.

#### Representational consistency at the image pixel level

To benchmark the representational space correlation for the input images, using pixel intensity as the input value, we averaged the pixel patterns for all images in a given category sharing one value of a nonidentity feature to generate a category pixel pattern (similar to how fMRI response patterns and CNN layer response patterns were generated for a given category). We calculated the Euclidean distances between pairs of category pixel patterns to generate an RDM for each value of a nonidentity feature and then correlated the RDMs between the two values of a nonidentity feature as we did before to assess the consistency in the pixel space across a given nonidentity feature change.

#### Comparing the brain and CNN responses

To directly compare between the brain and CNN results, for each of the three tolerance measures (i.e., rank-order preservation, cross-decoding success, and consistency), using Spearman rank correlation, we correlated the overall response profiles over the course of visual processing between a given CNN and each human participant. The resulting correlation coefficients were then tested against the lower bound of the noise ceiling of the brain data, with correlations below the lower bound indicating a given CNN measure to be significantly different from the corresponding measures from the human brain. When the number of layers sampled in a CNN did not match the number of brain regions tested, bilinear interpolation was used to down sample the CNN profile to match with that of the brain. This allowed us to preserve the overall response profile of the CNN while still being able to carry out the correlation analysis.

The upper and lower bounds of the noise ceiling were calculated following the procedure described in Nili et al. (2014). Specifically, the upper bound of the noise ceiling was calculated by taking the average of the Spearman correlation coefficients between each participant’s response and the group averaged response including all participants, whereas the lower bound of the noise ceiling was calculated by taking the average of the Spearman correlation coefficients between each participant’s response and the group average response excluding that participant.

To examine the coupling in response profiles among the three tolerance measures in CNNs and whether they showed the same coupling as those found in the brain, we also compared the pairwise correlation coefficients of the response profiles from these three measures between the brain and CNNs. Specifically, we evaluated whether or not the pairwise correlations obtained from the CNNs across all sampled layers were significantly outside the corresponding values obtained from the human brain.

#### Statistical analyses

Statistical analyses performed included direct comparisons of the obtained values with the baseline using t tests, correlations of the obtain measures with the rank order of the brain regions using Spearman rank correlation, and comparisons between the brain and CNN results as described above. One-tailed t test was used if a comparison was in a prespecified direction. Corrections for multiple comparisons using the Benjamini-Hochberg method (Benjamini & Hochberg, 1995) were made for the following number of comparisons: for the six brain regions when comparisons were performed within each nonidentity feature in the brain data, for the four types of nonidentity features when comparisons were assessed across the different nonidentity feature types in the brain data, and for the three types of comparisons made when comparing between the results of the brain and each CNN for each nonidentity feature change. When t tests were performed on correlation coefficients, values were Fisher-transformed first to ensure normal distribution.

As a correlation coefficient approached 1 or -1, because the corresponding Fisher-transformed values would approach the positive and negative infinities, respectively, for plotting purposes, any correlation coefficient greater than .95 or less than -.95 were replaced with the corresponding boundary value of .95 or -.95 before Fisher-transformation. To maintain consistency between the plots and the reported stats, statistical analyses were performed on these clipped data. Statistical results from the unclipped data were very similar to those from the clipped data.

## Acknowledgement

We thank Martin Schrimpf for help implementing CORnet-S, JohnMark Tayler for extracting the features from the three Resnet-50 models trained with the stylized images, and Brian Scholl and Nick Turk-Brown for helpful discussions and feedback on the results. This research was supported by National Institute of Health Grants (1R01EY030854 and 1R01EY022355) to Y.X. MVP was supported in part by NIH Intramural Research Program ZIA MH002035.

## Author contributions

The fMRI data used here were from two prior publications (Vaziri-Pashkam & Xu, 2019; Vaziri-Pashkam et al., 2019), with MV-P and YX designing the fMRI experiments and MV-P collecting and analyzing the fMRI data. YX conceptualized the present study and performed all the analyses reported here with input provided by MV-P. YX wrote the manuscript with comments from MV-P.

## Supplementary Results

### Decoupling of consistency measures at the image pixel level and those from the brain and CNNs

Could consistency in early visual areas be a direct reflection of consistency at the input image pixel level? To test this, using the same procedure (see Methods), we calculated consistency at the image level across nonidentity feature changes. Consistencies were 1.00, .92, .13 and -.33, respectively, for position, size, image stats and SF changes. The high consistencies at the image pixel level for position and size change were the result of the uniformity of the image space. However, these high consistencies were not reflected in brain responses, as consistencies in early visual areas for these two types of changes were quite low (i.e., around 0 for position and .25 for size change) and gradually increased from lower to higher visual areas. Likewise, despite the low consistencies obtained at the image level for image stats and SF changes, consistencies in early visual areas were much higher (i.e., around .75 for image stats and .4 for SF change). These results showed that consistency in early visual areas was not a direct reflection of consistency at the image pixel level.

In CNNs, for both position and size changes, high consistencies at the early layers were similar to those found at the image pixel level. However, for image stats change, consistency was low at the image pixel level (.13) but were much higher (>.5) in CNN early layers. Thus, high consistency at the image pixel level did not always translate to high consistency at early CNN layers either.

### No eye position confound

Could our results be due to a difference in eye position patterns for the different object categories across a nonidentity feature change? Such a difference could potentially result in fMRI response correlation for the different categories to differ across a nonidentity feature change, especially for early visual areas where the receptive field sizes are small. In other words, the eye positions for viewing the different categories could have followed one pattern at one value of a nonidentity feature and then followed a different pattern at the other value of the nonidentity feature.

For both the rank-order preservation and the consistency measures, we used the correlation between odd and even halves of the data within each value of the nonidentity feature as our reliability measure. We then normalized the correlation between the odd and even halves of the data across a feature change to this reliability measure. As such, even if eye positions for viewing the different categories were different, as long as such eye movement pattern across odd and even runs were similar with and without a nonidentity feature change, changes in eye positions for the different categories should have been accounted for. The only exception to this is when eye movement patterns across odd and even runs were more different with than without a feature change. For our decoding measures, because we used a leave-one-out cross-validation procedure by training the classifier on N-1 runs and then testing both the within- and cross-decoding on the Nth run, only when the averaged eye position for N-1 runs was more different from the Nth run with than that without a feature change, would eye position difference potentially contribute to a decoding difference.

All our participants had ample experience participating in prior fMRI experiments. In particular, they were selected based on their ability to maintain fixation during the long topographic mapping sessions to allow these maps in occipital and posterior parietal cortices to be reasonably well delineated. We see no reason why they should fail to maintain proper fixation in the present study. Nevertheless, to examine potential differences in eye position patterns, we analyzed the recorded eye position data.

To do so, we first removed saccades and blinks. We then corrected for eye position measurement drifts for each stimulus block in each run by subtracting the median deviation in eye position during the fixation block right before that stimulus block in that run (when only a single fixation dot was present on the screen). From the average X and Y eye positions for each stimulus block of each run, we either removed outliers that were 3 SD from the mean of all the data of a given participant in a given experiment or kept all the data (results did not differ whether or not outliers were removed). We analyzed the results separately for X and Y positions. We also concatenated the X and Y eye positions of each category together, separately for each value of the nonidentity feature for each half of the data. Each analysis contained four eye position pattern vectors (i.e., “odd1”, “odd2”, “even1”, and “even2”), each with N (for X or Y position) or 2N (for X and Y concatenated) unique categories x 2 feature values number of elements. For both the rank-order preservation and the consistency measures in which a split-half approach was used, to assess whether or not eye position patterns were more similar without than with a nonidentity feature change, we correlated the eye position pattern vector across the odd and even halves of the data in each participant. For the eye position pattern correlation without a feature change, we correlated “odd1” with “even1”, and “odd2” with “even2”, and took the average of the two correlation coefficients as the “same” correlation; and for the eye position pattern correlation across a feature change, we correlated “odd1” with “even2”, and “odd2” with “even1”, and took the average of the two correlation coefficients as the “different” correlation. We then Fisher z-transformed these correlation coefficients and tested at the participant group level whether “same” was different from “different”. That is, whether eye position patterns were more similar within than across a feature change across the two halves of the data. This was done for each experiment using paired t tests and the results were corrected for multiple comparisons for the four experiments included using the Benjamini-Hochberg method (Benjamini & Hochberg, 1995). To match the N-fold leave-one-out cross-validation procedure used for the decoding analysis, we repeated the above analysis but correlated eye position vectors from the average of N-1 runs with those from the Nth run, repeated the analysis with each run serving as the left-out run, and averaged the results.

No difference was found in eye position data in the split-half procedure in any of the four experiments whether or not outliers were included or excluded (without outliers, x positions, *ts* < 2.30, *ps* > .24; y positions, *ts* < 1.86, *ps* > .28; x and y positions concatenated, *ts* < 1.64, *ps* > .36; with outliers, x positions, *ts* < 2.74, *ps* > .13; y positions, *ts* < 1.46, *ps* > .36; x and y positions concatenated, *ts* < 1.66, *ps* > .59). Likewise, no difference was found in eye position data in the N-fold leave-one-out cross-validation procedure in any of the four experiments whether or not outliers were included or excluded (without outliers, x positions, *ts* < 1.66, *ps* > .50; y positions, *ts* < 1.07, *ps* > .57; x and y positions concatenated, *ts* < 2.60, *ps* > .11; with outliers, x positions, *ts* < 2.11, *ps* > .25; y positions, *ts* < .83, *ps* > .79; x and y positions concatenated, *ts* < 1.53, *ps* > .34).

We thus do not believe eye position confounds contributed to the present results.

## Supplementary Figure Captions

**Supplementary Figure 1.**
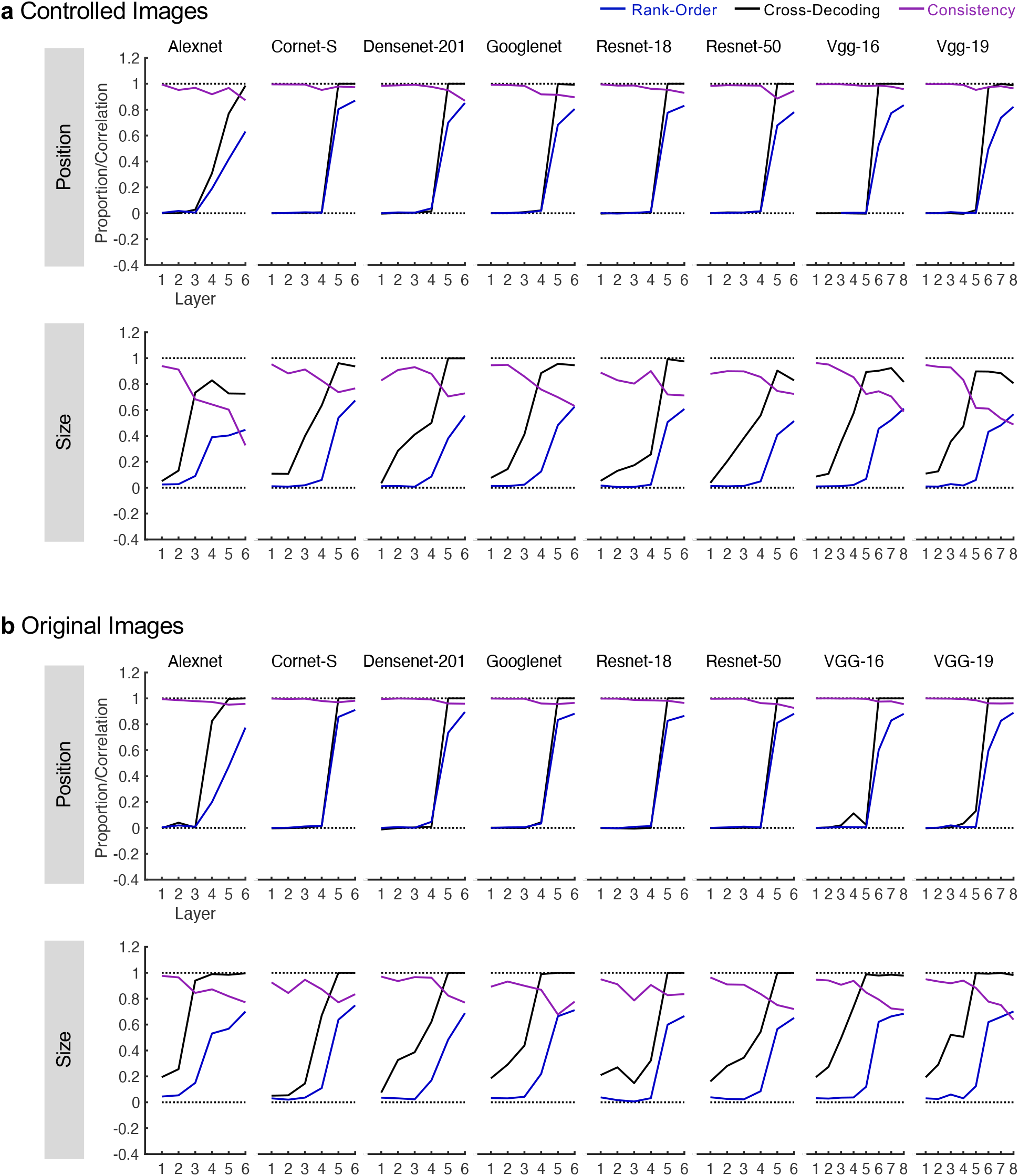
Rank-order preservation, cross-decoding success and consistency from 8 CNNs pretrained for image classification across the two Euclidean feature changes (position and size) using both the controlled and the original images. **a.** Results from the controlled images. These are the same results as those shown in Figure 4. **b.** Results from the original images. Very similar results were obtained with both types of images.

**Supplementary Figure 2.**
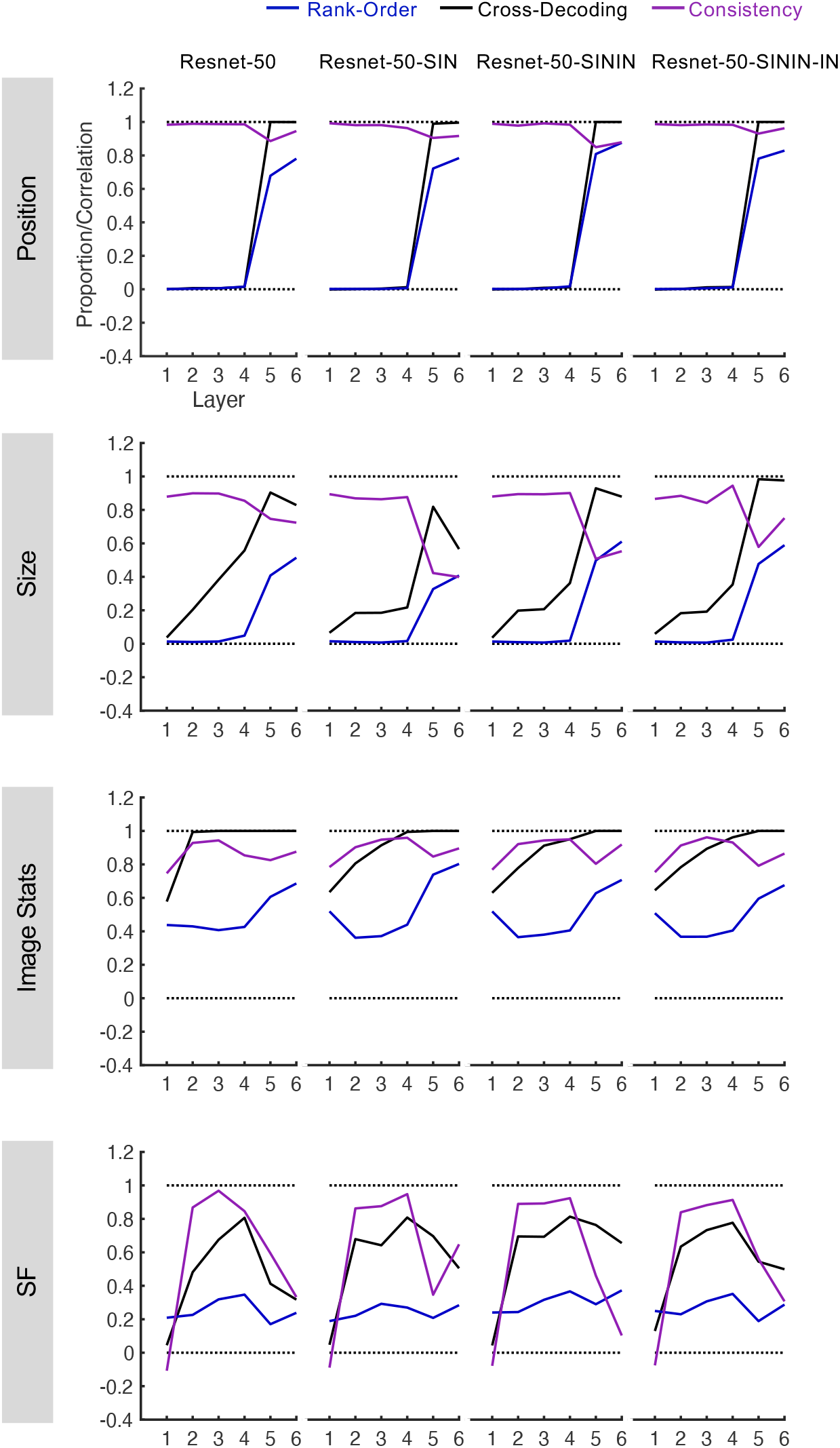
Rank-order preservation, cross-decoding success, and consistency from Resnet-50 with different training regimes across the two Euclidean (position and size) and the two non-Euclidean (image stats and SF) feature changes. Resnet-50 was pretrained either with the original ImageNet images (RN50-IN), the stylized ImageNet Images (RN50-SIN), both the original and the stylized ImageNet Images (RN50-SININ), or both sets of images and then fine-tuned with the stylized ImageNet images (RN50-SININ-IN).

**Supplementary Figure 3.**
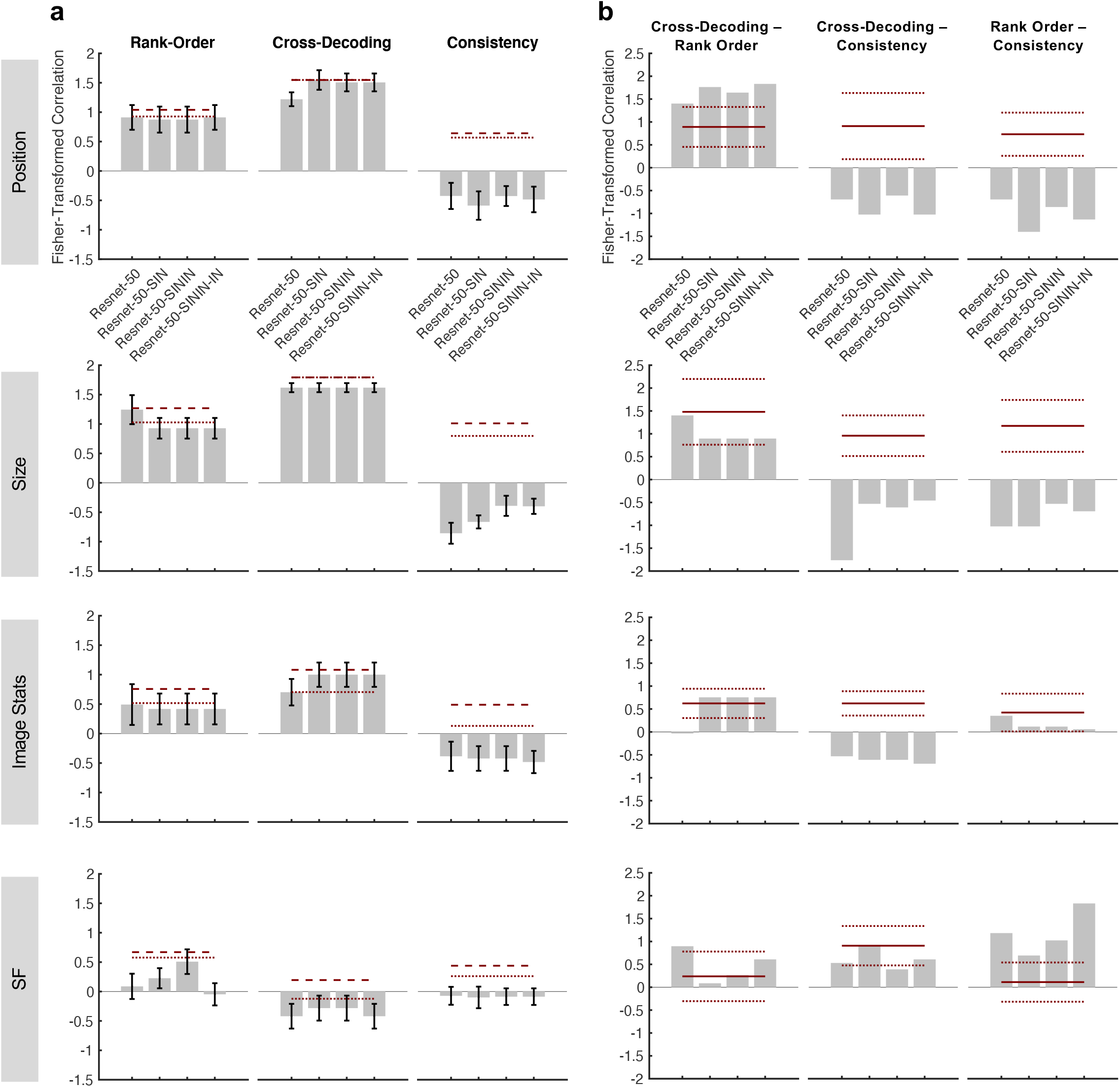
Correlations between the brain and Resnet-50 with different training regimes across the two Euclidean (position and size) and the two non-Euclidean (image stats and SF) feature changes. Resnet-50 was pretrained either with the original ImageNet images (RN50-IN), the stylized ImageNet Images (RN50-SIN), both the original and the stylized ImageNet Images (RN50-SININ), or both sets of images and then fine-tuned with the stylized ImageNet images (RN50-SININ-IN). **a.** Brain and CNN correlations of response profiles for rank-order preservation, cross-decoding success and consistency across the two Euclidean and the two non-Euclidean nonidentity feature changes. The dashed and dotted dark red lines represent the upper and lower bounds of the noise ceiling, respectively, calculated from the brain data. **b.** Brain and CNN comparisons of the coupling in response profiles among rank-order preservation, cross-decoding success and consistency for the two Euclidean and the two non-Euclidean nonidentity feature changes. The solid dark red lines represent the corresponding mean correlation measures from the brain, with the dotted dark red lines indicating the 95% confidence intervals around the means. All values were Fisher z-transformed to ensure valid statistical comparisons.

**Supplementary Figure 4.**
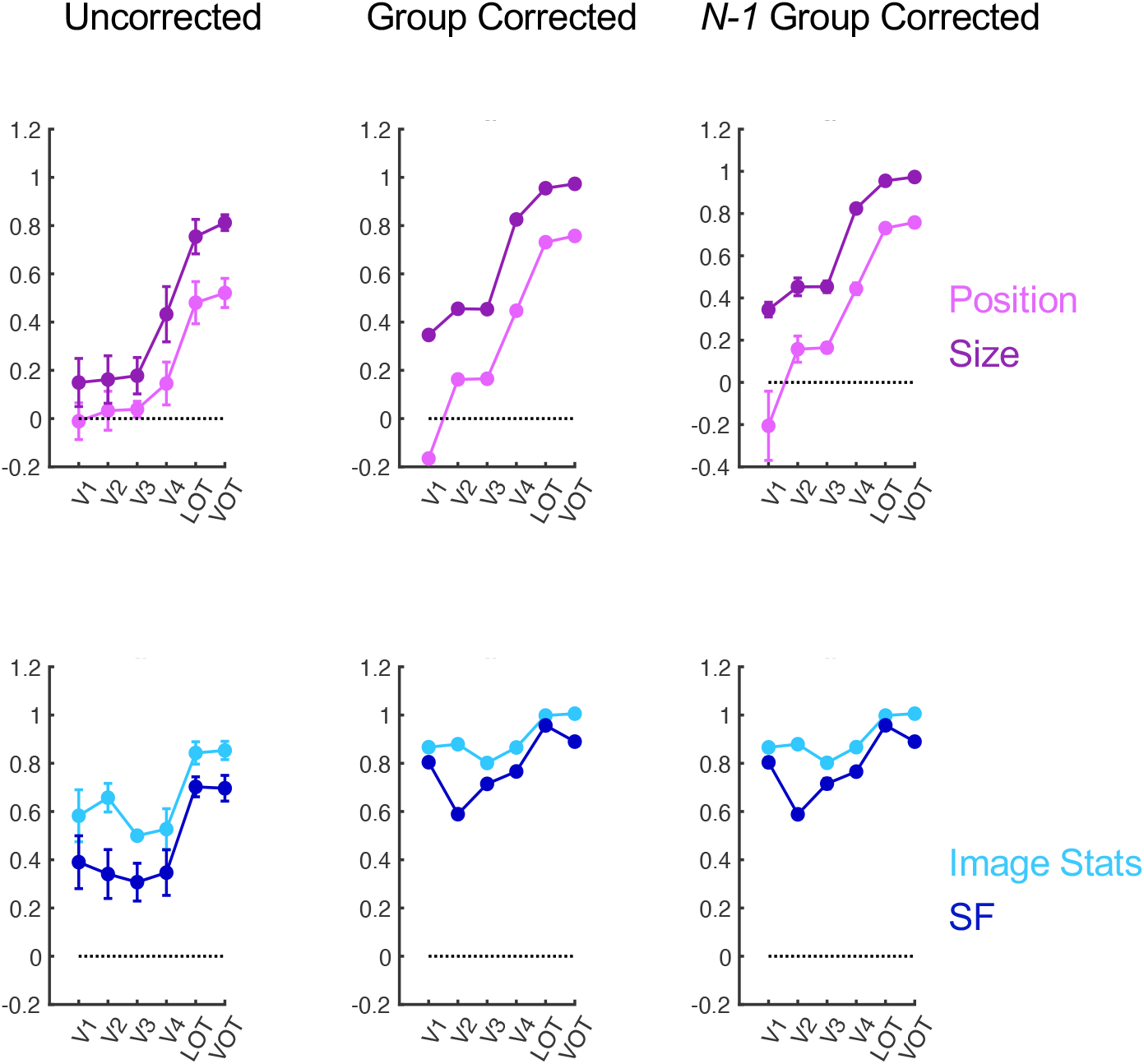
Evaluating representational space consistency across the two Euclidean (position and size) and the two non-Euclidean (image stats and SF) feature changes in the human brain with different reliability correction methods. Left column, Uncorrected. Here no reliability correction was applied. Middle column, Group Corrected. Here both correlation and reliability measures were first averaged across all participants. Reliability correction was then applied to these averaged values. Right column, *N-1* Group Corrected. Here group corrected correlations from *N-1* participants were computed first and then averaged across all *N* iterations. Error bars represent standard errors of the means.

